# Prevention of *Unc13a* cryptic splicing is sufficient to preserve memory

**DOI:** 10.64898/2026.09.11.751031

**Authors:** Tianyu Cao, Meghraj Singh Baghel, Rashmi Thapa, Sishir Gautam, Aswathy Peethambaran Mallika, Yijun Wei, Xiaoke K Chen, Irika R. Sinha, Grace D. Burns, Shruti Renganathan, Xinrui Wen, Bo Pang, Jihee Choi, Jonathan P Ling, Da-Ting Lin, Rongsong Liu, Yun Li, Philip C. Wong

## Abstract

TDP-43 dysfunction is thought to underlie frontotemporal dementia and limbic-predominant age-related TDP-43 encephalopathy, neurodegenerative dementias currently without effective therapy. Therapeutic strategies are designed to correct individual cryptic targets of TDP-43, such as *UNC13A,* whereby its cryptic splicing compromises synaptic function, yet the sufficiency of such an approach to prevent memory deficits is unclear. Using a forebrain neuron-specific TDP-43 knockout mouse model that recapitulates TDP-43 dysfunction occurring during early stages of human disorders, we found here that prevention of cryptic splicing to include that of *Unc13a* attenuated memory deficits. We show that genetic ablation of *Unc13a* cryptic exon solely in such TDP-43 knockout mice is sufficient to preserve cognition, supporting the clinical value of targeting *UNC13A* to mitigate memory deficits. Prevention of cryptic splicing of multiple targets of TDP-43 additionally attenuate neuron loss. For optimal outcomes in TDP-43 related dementias, these findings thus strongly support strategies designed to repress cryptic splicing of multiple targets of TDP-43, including *UNC13A*.

## Introduction

Loss of nuclear TAR DNA/RNA-binding protein 43kDa (*TARDBP*, TDP-43) and its splicing repression is recognized as a central pathogenic mechanism across a broad spectrum of adult-onset neurodegenerative diseases^1–3^, including frontotemporal lobar degeneration with TDP-43 pathology (FTLD-TDP), amyotrophic lateral sclerosis (ALS), limbic-predominant age-related TDP-43 encephalopathy (LATE), Alzheimer’s disease with LATE (AD-LATE), devastating disorders^4–14^ currently without disease-modifying therapy. TDP-43, a highly conserved nuclear RNA-binding protein, was initially implicated in disease through its cytoplasmic aggregation^15^ however, a major function of TDP-43 was subsequently shown to regulate splicing repression of cryptic exons, the loss of which is a key pathogenic feature of TDP-43 proteinopathies^1^. This discovery led to the proposal that deficits in TDP-43-dependent cryptic splicing represent an early pathogenic event that drives neuron loss, occurring independently of, and prior to, the cytoplasmic aggregation of TDP-43. Several lines of evidence support this view. First, cryptic exons encoded peptides, such as that found in cryptic *HDGFL2*, can be found in biofluids at presymptomatic stage of *C9orf72* carrier ALS-FTD patients^2,3^ and TDP-43 splicing dysfunction preceding its cytoplasmic aggregates in the human aging brain by at least a decade^16^. Antemortem brain tissue from a presymptomatic *C9orf72* carrier^17^ and postmortem analyses of FTLD-TDP^18^ and AD^19^ brains reveal nuclear depletion of TDP-43 and cryptic exon inclusion in neurons that lack cytoplasmic aggregates. Second, ALS patient-derived iPSC neurons revealed loss of TDP-43 splicing repression even in the absence of its aggregation^20^. Third, mutations in TDP-43 linked to ALS^21–23^ impact a TDP-43 cryptic exon of stathmin-2 (*STMN2*) in human iPSC-derived neurons, independent of TDP-43 cytoplasmic aggregates^24–26^. Furthermore, *UNC13A* encoding a presynaptic protein critical for neurotransmitter release^27–30^, harbors strong risk alleles for ALS-FTD, which influences the inclusion of its cryptic exon^31,32^. Antisense oligonucleotides (ASOs) designed to correct *STMN2* or *UNC13A* cryptic splicing have entered clinical testing for ALS^33–35^. While these strategies targeting individual cryptic exon of TDP-43 hold promise, it is not clear whether prevention of any one cryptic target, such as *UNC13A,* is sufficient to preserve cognition.

To address this question, we used our conditional knockout mouse model lacking TDP-43 in forebrain neurons (*CamKIIa-CreER;Tardbp^f/f^*, (hereafter referred to as TDP-43cKO) mimicking the early TDP-43 dysfunction observed in biofluids of ALS-FTD patients^2^. We previously showed that loss of TDP-43-dependent cryptic splicing leads to forebrain circuit abnormalities^36^, memory deficits and brain atrophy^37,38^. To restore TDP-43 splicing repression in adult forebrain neurons, we employed a previously characterized splicing repressor, termed CTR^1^, which retains TDP-43’s RNA recognition domains while replacing its aggregation-prone low-complexity domain with the splicing repression domain of RAVER1^39–41^. The construct further incorporates the human *TARDBP* 3′UTR to preserve autoregulatory feedback, enabled splicing repression of many cryptic targets^1,42,43^. CTR was delivered intracerebroventricularly to adult TDP-43cKO mice utilizing an AAV-PHP.eB^44,45^ that facilitates efficient transduction of adult central neurons. AAV-PHP.eB-CTR restored splicing repression across many TDP-43-regulated targets and rescued cognitive and circuit deficits in TDP-43cKO mice. To then test the sufficiency of correcting a single, translationally relevant cryptic target to rescue cognitive deficits and neuron loss, we focus on cryptic *Unc13a*, an unexpected, shared target between humans and mice; while the human cryptic exon (localized within intron 20) is distinct from that of the mouse (intron 1), they both introduce a premature termination codon leading the nonsense mediated decay of the mRNA (**Fig. 1d**) and is the only such target that permits direct genetic dissection *in vivo*. We genetically deleted the *Unc13a* cryptic exon in TDP-43cKO mice and evaluated whether preventing inclusion of *Unc13a* cryptic exon alone was sufficient to rescue cognitive deficits^1,42,43^.

**Figure 1.**
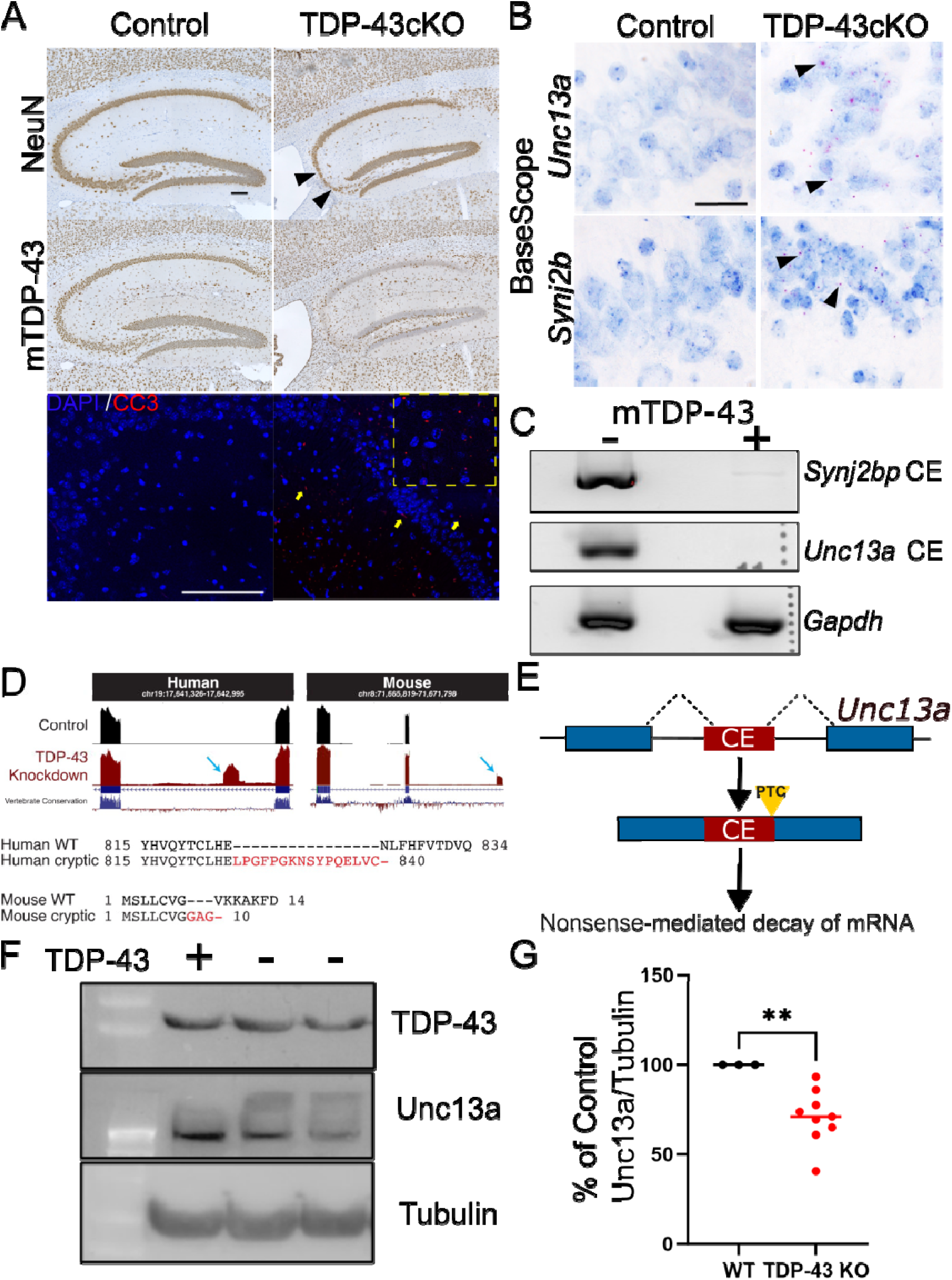
Inclusion of *Unc13a* cryptic exon reduced Unc13A protein in TDP-43cKO mice. (**A**) Immunohistochemical analysis using mouse-TDP-43 and NeuN antisera in brains of TDP-43cKO and control mice (arrow heads indicate degenerated neurons, scale bar 200µm). Representative images of immunofluorescence staining of cleaved caspase 3 (CC3) in control and TDP-43cKO mice (yellow arrowheads indicate CC3 positive puncta in hippocampus with inset view, scale bar 100µm). (**B**) BaseScope *in-situ* RNA hybridization analysis of *Unc13a* (upper panel) and *Synj2bp* (lower panel) CEs in control and TDP-43cKO mice (arrowheads indicate pink signal of cryptic RNA foci, scale bar, 40µm). (**C**) RT-PCR analysis of *Unc13a* and *Synj2bp* cryptic exon in control and TDP-43cKO mice. (**D**) Schematic representation of inclusion of *Unc13a CE* as a common shared target gene of TDP-43 in human and mouse data sets (arrow indicates cryptic exon inclusion in human and mouse transcriptome). (**E**) Schematic representation of *Unc13a* CE inclusion which introduces <u>P</u>remature <u>T</u>ermination <u>C</u>odon (PTC) leading to non-sense mediated decay of mRNA. (**F, G**) Immunoblotting analysis of TDP-43 and Unc13A in littermate control (n=3) and TDP-43cKO (n=9) mice, correlating loss of TDP-43 to reduction of level of Unc13A (sample t-test; **P<0.01).

## Results

### *Unc13a* cryptic exon inclusion reduces its protein level in TDP-43cKO mice

To model the early stage of TDP-43 dysfunction occurring in human disease, we selectively deleted TDP-43 from forebrain excitatory neurons of adult TDP-43cKO mice. Initially, we confirmed the depletion of TDP-43 from neuronal nuclei of the hippocampus (Fig. 1A, middle panel) and cortex (data not shown) of TDP-43cKO mice, while control littermates (Tardbp^f/f^) maintained normal TDP-43 level as expected. Loss of TDP-43 was accompanied by activation of cleaved caspase-3 (CC3) and progressive neuron loss in CA2/3 neurons of the hippocampus (**Fig. 1A, top and lower panels**). Upon TDP-43 depletion, we observed *Unc13a* and *Synj2bp* cryptic RNA in TDP-43-deficient forebrain neurons by RNA *in situ* hybridization (**Fig. 1B**) or RT-PCR analysis (**Fig. 1C**). Among TDP-43-regulated cryptic exons, *UNC13A* is unique in that while it is a shared cryptic target between humans and mice albeit their cryptic exons are localized within different intronic regions (**Fig. 1D**). Human studies have shown that inclusion of *UNC13A* cryptic exon introduced premature termination codon (PTC), leading to nonsense-mediated decay of *UNC13A* transcripts and reduction of UNC13A protein ^31,32,46^. Bioinformatics analysis predicted that *Unc13a* cryptic exon also introduces PTC in mice, which should also lead to nonsense-mediated decay of *Unc13a* transcripts and reduction of Unc13a protein (**Fig. 1E**). We confirmed this prediction by showing a marked reduction of Unc13a in hippocampus of TDP-43cKO mice (**Fig. 1F, G**). Together, these data establish that loss of TDP-43 in forebrain excitatory neurons recapitulates a key feature of TDP-43 pathology, including cryptic splicing of a conserved target such as *Unc13a*, reduction of Unc13a protein, caspase-3 activation, and selective hippocampal neuron loss.

### AAV-PHP.eB-CTR restores TDP-43 splicing repression in excitatory neurons lacking TDP-43

Having established that TDP-43 depletion drives cryptic splicing and neurodegeneration in forebrain excitatory neurons, we tested whether transcriptome-wide restoration of TDP-43 splicing repression could mitigate these abnormalities. To validate such ability of CTR (**Sup Fig. 1C**) to rescue cell death of forebrain neurons lacking TDP-43, we delivered AAV9-CTR intracerebroventricularly (ICV) to P0 *CamKIIa-CreER;Tardbp^f/f^* pups. We found that AAV9-CTR attenuates neuronal loss occurring in CA2/3 hippocampal region neurons of mice lacking TDP-43 (**Sup Fig 2A-C**). Corroborating these results, we showed that TDP-43 cryptic exons are repressed in rescued mice (**Sup Fig. 2D-F**), confirming that the failure to repress TDP-43 cryptic exons in central neurons underlies neuronal loss. However, for clinically relevant context, it is critical to establish delivery of CTR using an AAV serotype that broadly transduces central neurons to restore TDP-43 dysfunction in the adult brain.

For efficient CTR delivery in adult forebrain neurons of *CamKIIa-CreER;Tardbp^f/f^* mice, we delivered the chimeric splicing repressor CTR to adult TDP-43cKO mice by intracerebroventricular (ICV) injection of AAV-PHP.eB, a BBB-crossing serotype that facilitates efficient transduction of adult central neurons^44,45^. Tamoxifen was administered at 5 months of age to deplete TDP-43, followed by ICV injection of AAV-PHP.eB-CTR or AAV-PHP.eB-GFP control 2 weeks later (**Sup Fig. 1D**). Immunohistochemical analysis using an antibody specific to the N-terminal region of human TDP-43 (which detects CTR but not endogenous mouse TDP-43), revealed widespread CTR expression throughout the hippocampus, with robust signal in neurons of CA2/3 and detectable expression in those of CA1 **(Fig. 2A, second and third column)**. Quantification of transduction efficiency showed an average rate of ∼40% of neurons across 1, 3, and 6 months post-injection (**Fig. 2B**), indicating durable transgene expression.

**Figure 2.**
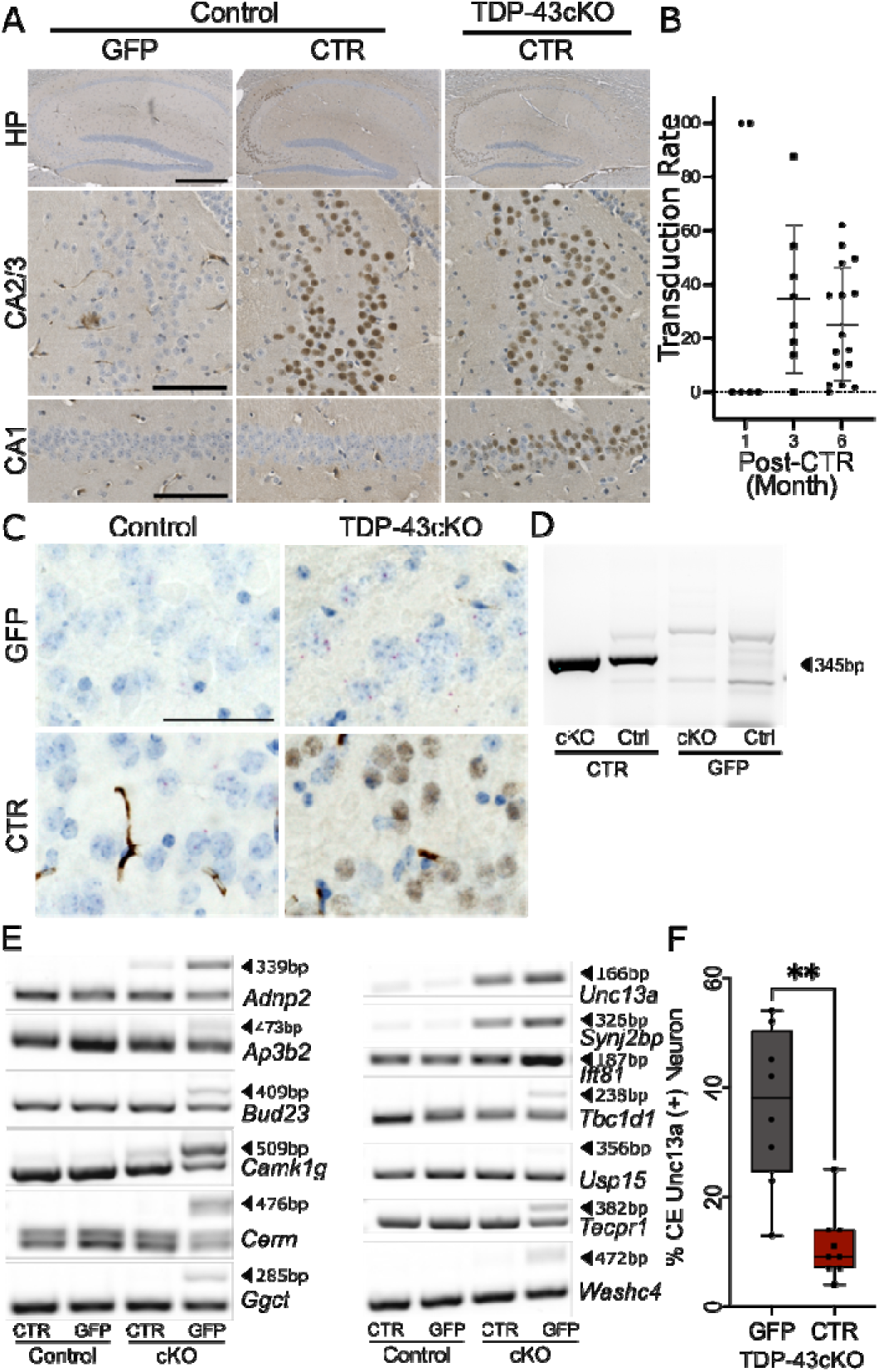
AAV-PHP.eB-CTR restores TDP-43 splicing repression in excitatory neurons lacking TDP-43. (**A**) Representative immunohistochemistry showing widespread CTR protein expression in the hippocampus of TDP-43cKO and control mice following ICV of AAV-CTR. CTR expression was detected using an antibody targeting the human TDP-43 N-terminus. Strong expression was observed in the CA2 and CA3 subregions, with additional signal in CA1. (**B**) Quantification of CTR viral transduction rate in TDP-43cKO mice at 1-, 3-, and 6-months post-injection(n = 6,8,16, respectively). Transduction rates remained stable over time (∼40%). (**C**) Representative images of BaseScope *in situ* hybridization & Immunohistochemistry co-detection for cryptic *Unc13a* RNA and CTR fusion protein reveal the spatial distribution of splicing defects in hippocampus CA2/3 subregions. CTR protein, which marked by N-terminal antibody of human TDP-43 protein, showed an expression in treated TDP-43cKO mice, and less expressed in treated control mice. Cryptic *Unc13a* transcripts were elevated in dentate gyrus (DG), CA1, and CA2/3 of TDP-43cKO-GFP mice and markedly reduced in CTR-treated animals, indicative of CTR restoring TDP-43 splicing regulation. (**D**) RT-PCR quantification of CTR mRNA levels using primers spanning the TDP-43 N-terminal and RAVER1 C-terminal sequences. Despite equal AAV dosing, CTR transcript levels were significantly higher in TDP-43cKO mice than in controls, consistent with autoregulatory stabilization of expression via the Tardbp 3′UTR. (**E**), Representative RT-PCR analysis of hippocampal RNA showing repression of multiple cryptic exon–containing transcripts in CTR-treated TDP-43cKO mice. Representative targets include *Adnp2, Ap3b2, Bud23, Camk1g, Cerm, Ggct, Unc13a, Synj2bp, Tbc1d1, Usp15, Tecpr1,* and *Washc4*. (**F**) Quantification of *Unc13a* cryptic exon inclusion percentage relative to total neuron count. TDP-43cKO mice treated with GFP (n=8) exhibited a significant increase in cryptic exon usage compared to CTR-treated cKO (n=9, p=0.0011). Scale bars: A: upper panel 500 μm, lower panel 100 μm.; C: Error bars represent mean ± s.e.m. P-values were calculated using two-tailed unpaired t-tests or Mann–Whitney tests as specified in figure panels; n values are defined in the corresponding Methods.

We next assessed whether CTR expression was sufficient to restore TDP-43’s splicing repressor function *in vivo*. RNA *in situ* hybridization revealed that cryptic *Unc13a* RNA accumulated in multiple hippocampal subregions upon TDP-43 depletion, with the highest signal in the dentate gyrus, followed by CA1 and CA2/3 (**Fig. 2C, upper panel; Sup Fig. 3**). CTR treatment markedly reduced cryptic *Unc13a* expression across all subregions as compared to GFP-treated TDP-43cKO mice, as assessed by co-detection of CTR protein with cryptic *Unc13a* RNA (**Fig. 2C, lower panel second column; Sup Fig. 3**). Importantly, RT-PCR analysis demonstrated that CTR attenuated cryptic splicing not only of *Unc13a* but across a panel of TDP-43-regulated cryptic transcripts, including *Adnp2, Ap3b2, Bud23, Camk1g, Crem, Ggct, Synj2bp, Tbc1d1, Usp15, Tecpr1,* and *Washc4* (**Fig. 2E**). As compared to GFP-treated mice, the levels of cryptic exon inclusion for *Unc13a* in CTR-treated TDP-43cKO mice were significantly lower (**Fig. 2F**). Together, these data indicate that CTR functionally restored multiple cryptic targets of TDP-43 in the adult brain.

A critical design feature of CTR is the inclusion of the human *TARDBP* 3′UTR, which preserves TDP-43’s autoregulatory feedback mechanism^47,48^ (**Sup Fig. 1C**). To confirm that this element functions *in vivo*, we quantified CTR mRNA levels using RT-PCR with primers spanning the TDP-43 and RAVER1 junction. Despite equivalent AAV doses, CTR transcript levels were significantly higher in TDP-43cKO mice than that of controls (**Fig. 2D**), consistent with autoregulation of CTR expression to within physiological levels. This result confirms that CTR expression is upregulated when it is needed and constrained when endogenous TDP-43 is present, a property essential for preventing overexpression toxicity within a gene therapy context^49^.

### AAV-PHP.eB-CTR normalizes circuit activity and rescues cognitive deficits

To determine whether the restoration of splicing repression by CTR translates into functional recovery, AAV-PHP.eB-CTR treated mice underwent a battery of behavioral assessments. No significant differences in locomotor activity or anxiety-related behavior were observed between genotypes or treatment groups **(Sup Fig. 4)**. However, in the novel object recognition test, TDP-43cKO mice treated with AAV-PHP.eB-RFP as control failed to discriminate between familiar and novel objects, whereas AAV-PHP.eB-CTR treated TDP-43cKO mice displayed restored novelty preference comparable to control littermates **(Fig. 3A, D)**. Similarly, in the social behavior test, control groups treated with AAV-PHP.eB-RFP or AAV-PHP.eB-CTR displayed normal social behavior, but AAV-PHP.eB-RFP treated TDP-43cKO mice failed to show preference for a novel conspecific over a familiar one showing deficit in social behavior, and this deficit was fully rescued by CTR treatment **(Fig. 3B, E)**. These results demonstrate that CTR rescues both memory and social behavior deficits caused by TDP-43 dysfunction.

**Figure 3.**
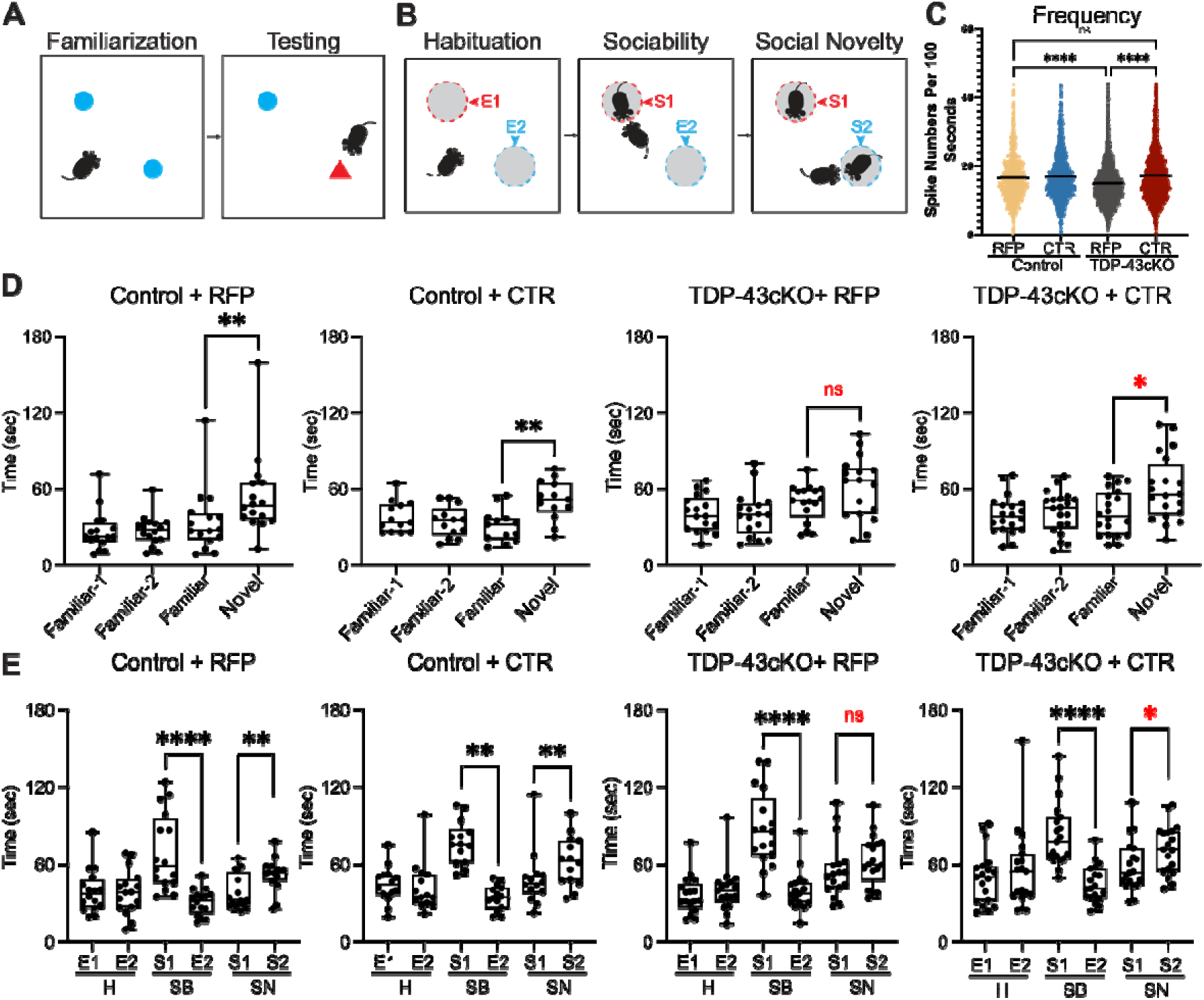
AAV-PHP.eB-CTR restores neuronal calcium activity and attenuates memory deficits. (**A**) Schematic of the novel object recognition (NOR) test protocol. Mice were exposed to two identical objects during a familiarization phase, followed by a testing phase in which one object was replaced with a novel item. The time investigating each object was measured. (**B**) Schematic of the social behavior test. Mice underwent three 10-minute phases: habituation (two empty cages), sociability (one unfamiliar mouse introduced), and social novelty (second unfamiliar mouse added). (**C**) Analysis of *in vivo* neuronal activity using calcium imaging in the prelimbic cortex. Compared to Control-RFP controls, TDP-43cKO-RFP mice exhibited reduced neuronal calcium spike frequency, consistent with impaired excitatory activity. CTR treatment normalized activity levels. Imaging was performed through GRIN lenses following AAV-GCaMP6f injection. Group sizes: Control-RFP (n = 936 ROIs), Control-CTR (n = 1379 ROIs), TDP-43cKO-RFP (n = 2285 ROIs), TDP-43cKO-CTR (n = 2334 ROIs). **** indicates P < 0.0001, ns = not significant. (**D**), Quantification of exploration time during the novel object recognition (NOR) test. TDP-43cKO-RFP mice (n = 17) failed to show a preference for the novel object, indicating impaired recognition memory. CTR treatment (n = 20) restored novel object preference to levels comparable to Control-RFP (n = 16) and Control-CTR (n = 13) groups. (**E**), Quantification of time spent investigating social targets. While all groups displayed normal sociability, TDP-43cKO-RFP mice failed to show a preference for social novelty. CTR-treated TDP-43cKO mice showed restored preference for the novel conspecific, indicating rescue of social memory. Data are mean ± s.e.m. P-values were calculated using Kruskal–Wallis ANOVA followed by Dunn’s post hoc tests or Mann–Whitney tests, as noted in Methods. ****P<0.0001, **P<0.01, *P<0.05, ns = not significant.

To assess whether behavioral rescue reflected restoration of underlying neural circuit activity, we performed *in vivo* calcium imaging in the prelimbic cortex using a miniature fluorescence microscopy. AAV1-CamKII-GCaM P6f was stereotaxically injected into the prelimbic cortex, and a gradient-index (GRIN) lens was implanted at the injection site to enable single-cell resolution recording in freely behaving mice. TDP-43cKO mice treated with AAV-PHP.eB-RFP showed significantly reduced calcium transient frequency as compared to that of control littermates, indicating impaired excitatory neuronal activity (**Fig. 3C**). Importantly, AAV-PHP.eB-CTR treatment restored firing frequency to levels comparable to those of control mice (**Fig. 3C**), indicating that CTR rescues prefrontal neural circuit activity, in addition to memory deficits.

### Ablation of *Unc13a* cryptic exon is sufficient to preserve cognition in TDP-43cKO mice

Having established that restoration of multiple TDP-43 cryptic targets by CTR rescues cognitive deficits, we asked whether correction of a single cryptic target is sufficient to achieve the same outcome. To directly test whether preventing *Unc13a* cryptic exon inclusion is sufficient to mitigate disease phenotypes, we used CRISPR-Cas9 to delete a ∼44 bp region (**Fig. 4A**) from the mouse genome that defined the *Unc13a* cryptic exon (termed *Unc13aCE^-/-^* mice). Targeted deletion was confirmed by PCR and DNA sequencing analysis (**Sup Fig. 5A-B**), and the level of Unc13a in *Unc13aCE*^−/−^ mice were indistinguishable from that of wild-type mice (**Sup Fig. 6A**), confirming that deletion of *Unc13a* cryptic exon does not disrupt normal Unc13a expression.

**Figure 4.**
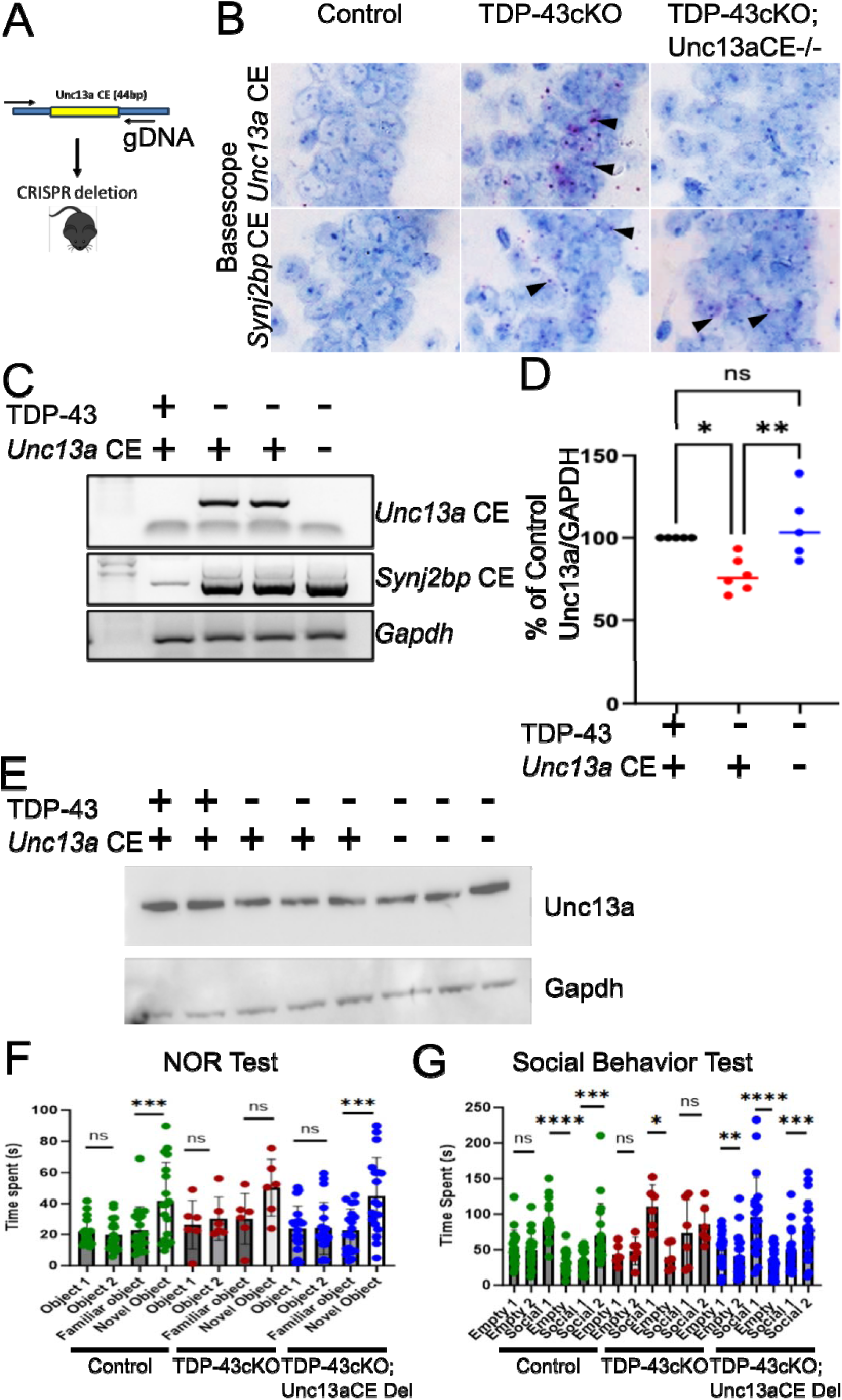
Restoring physiological level of Unc13A protein rescues cognitive deficits. (**A**) Schematic representation of *Unc13a CE* sequence (44bps) deletion by CRISPR technology, forward and reverse arrows represents the primers used for genotyping using PCR amplification. (**B**) Representative images of RNA in-situ analysis by BaseScope of *Unc13a* and *Synj2bp* in control (WT), TDP-43 deficient (TDP-43cKO) and TDP-43cKO;Unc CE^-/-^ mice (arrowheads indicate pink signal of cryptic RNA foci). (**C**) RT-PCR analysis of *Unc13a* and *Synj2bp* cryptic exon in control, TDP-43cKO and TDP-43cKO;Unc CE^-/-^ mice, showing deletion of Unc13a CE sequence prevented inclusion of *Unc13a* CE but not *Synj2bp* showing TDP-43 deletion. (**D, E**) Immunoblotting analysis of Unc13A protein in wild-type control (n=5), TDP-43 deleted (n=6) and TDP-43cKO;Unc CE^-/-^ (n=5) mice, quantification showing Unc13a CE deletion restores the Unc13A protein in TDP-43cKO;Unc CE^-/-^ mice (one-way ANOVA; ns: no significant difference; *P<0.05; **P<0.01). (**F**) Quantification of exploration time during the novel object recognition (NOR) test. TDP-43cKO mice (n=6) failed to show a preference for the novel object, indicating impaired recognition memory. Unc13a CE deletion (n = 16, included heterozygous and homozygous *Unc13a* CE deleted mice) restored novel object preference to levels comparable to control (n = 17) groups. (**G**) Quantification of time spent investigating social targets. While all groups displayed normal sociability, TDP-43cKO mice failed to show a preference for social novelty. TDP-43cKO;Unc CE^-/-^ mice showed restored preference for the novel conspecific, indicating rescue of social memory. P-values were calculated using Kruskal–Wallis ANOVA followed by Dunn’s post hoc tests or Mann–Whitney tests, as noted in Methods. (****P<0.0001, **P<0.01, *P<0.05, ns = not significant).

We then used *UncC13aCE*^−/−^ and TDP-43cKO mice in a crossbreeding strategy (**Sup Fig. 6B**) to generate compound mutants lacking TDP-43 in forebrain neurons and the *Unc13a* cryptic exon (TDP-43cKO;*Unc13aCE^-^*^/-^). Following tamoxifen-induced TDP-43 deletion at 2 months of age, RT-PCR and RNA *in situ* analysis at 4 months post-excision confirmed the absence of cryptic *Unc13a* transcripts in forebrain neurons of TDP-43cKO;*Unc13aCE*^-/-^ mice (**Fig. 4B, upper panel; C**); Importantly, cryptic exon inclusion of *Synj2bp*, another TDP-43 target, persisted in these animals (**Fig. 2B, lower panel; C),** confirming that TDP-43 cryptic splicing is maintained as expected except for that of *Unc13a*. Consistent with the suppression of nonsense-mediated decay, the level of Unc13a was normalized in TDP-43cKO;*Unc13aCE*^−/−^ mice (**Fig. 4D-E**). To determine whether normalizing the level of Unc13a could rescue the cognitive and social behavioral deficits caused by TDP-43 dysfunction, we first assessed novel object recognition tests. TDP-43cKO mice failed to discriminate between familiar and novel objects, whereas TDP-43cKO;*Unc13aCE*-/- mice displayed restored novelty preference comparable to that of controls (**Fig. 4F**). A similar rescue was observed in the social behavior test: TDP-43cKO;*Unc13aCE-/-* mice showed significant preference for a novel conspecific over a familiar one, in contrast to TDP-43cKO mice, which showed no such preference (**Fig. 4G**). These results demonstrate that preventing *Unc13a* cryptic splicing alone is sufficient to preserve memory, suggesting that *UNC13A* cryptic splicing is a key driver of cognitive deficit downstream of TDP-43 dysfunction and ASO designed to block *UNC13A* cryptic splicing hold promise for dementias associated with TDP-43 proteinopathy.

### AAV-PHP.eB-CTR mitigates loss of hippocampal neurons and attenuates brain atrophy

Having shown that both AAV-PHP.eB-CTR and *Unc13a* cryptic exon deletion preserve cognition in TDP-43-deficient mice, we examined whether either intervention attenuates neuron loss. Quantification of neuron numbers in the hippocampus of mice at 12 months post-injection revealed severe neuron loss in the CA2/3 subregion of TDP-43cKO mice exposed to AAV-PHP.eB-GFP (**Fig. 5A-B, second column**), as expected³ ⁻³. AAV-PHP.eB-CTR treated TDP-43cKO mice retained substantially more NeuN-positive neurons in CA2/3 (**Fig. 5A-C, third column**). The degree of neuroprotection closely matched the local transduction rate of approximately 40%, supporting a direct relationship between CTR expression and neuronal survival (**Fig. 5B**). This neuroprotective effect was durable: partial rescue was already evident at 6 months and became statistically significant at 12 months (**Fig. 5B-C**, **Sup Fig. 7**), indicating sustained protection across disease progression. As compared to AAV-PHP.eB-GFP, morphometric analysis further confirmed a significant rescue of hippocampal area in AAV-PHP.eB-CTR treated TDP-43cKO mice (**Fig. 5D; Sup Fig. 8**). Caspase-3 activation was markedly reduced in AAV-PHP.eB-CTR treated TDP-43cKO mice to levels indistinguishable from littermate controls (**Sup Fig. 9**). Importantly, long-term CTR exposure in *Tardbp^f/f^*mice, in which endogenous TDP-43 is normal, did not reduce hippocampal neuron number or area, nor did it produce overt adverse phenotypes (**Fig. 5A, first column; D**), consistent with the autoregulatory constraint conferred by the *TARDBP* 3′UTR (**Fig. 2D**).

**Figure 5.**
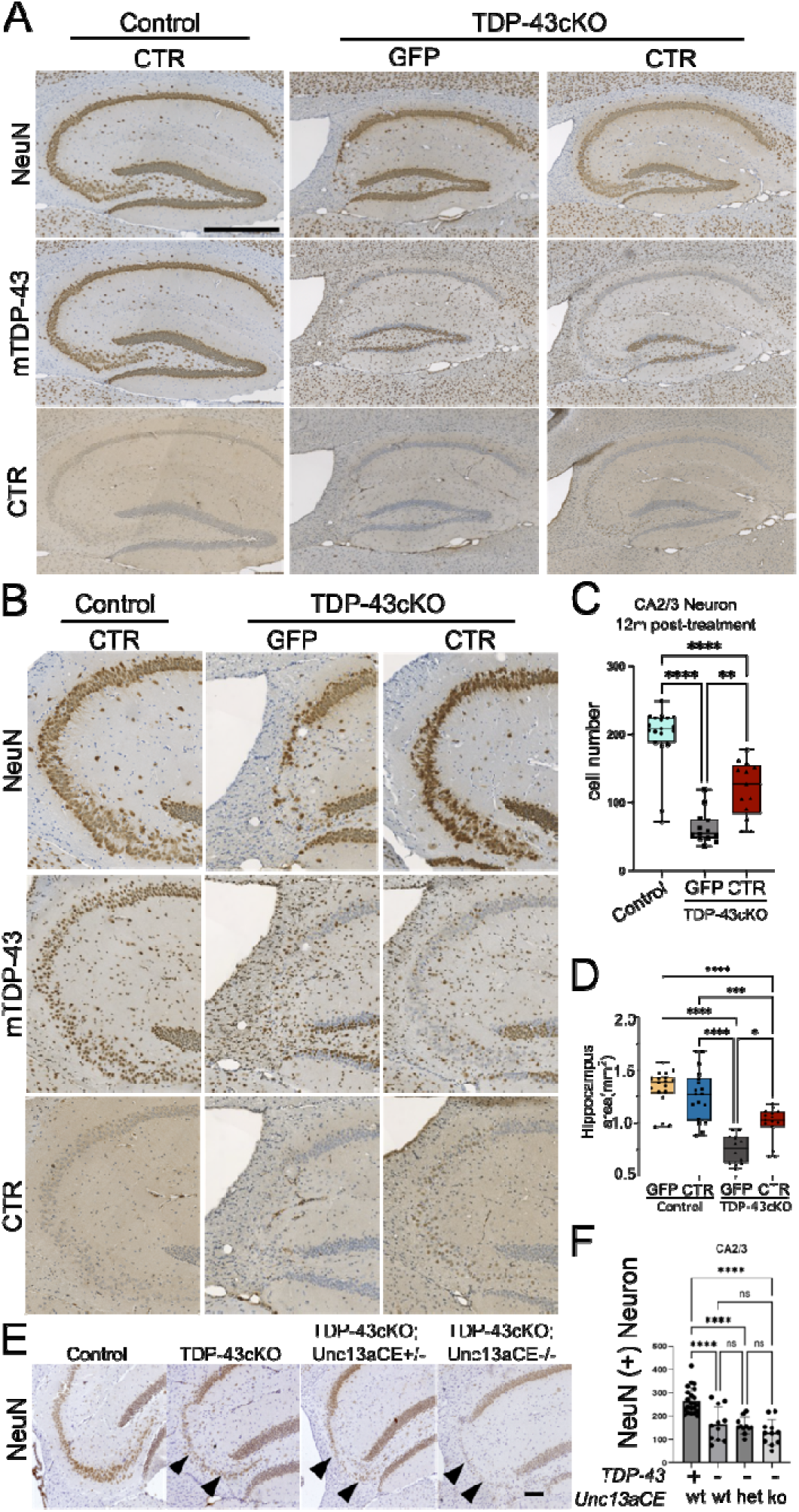
AAV-PHP.eB-CTR mitigates selective loss of hippocampal CA3/2 neurons and brain atrophy. (**A**) NeuN immunostaining overview of the entire hippocampus showing global anatomical differences across groups. CTR-treated TDP-43cKO mice exhibited visibly preserved hippocampal structure. (**B**) Representative NeuN immunohistochemistry images of the CA2/3 hippocampal subregion in control and TDP-43cKO mice 12 months after AAV-CTR or AAV-GFP injection. TDP-43cKO-GFP mice showed severe neuronal loss in this region, whereas TDP-43cKO-CTR mice retained more NeuN-positive cells. (**C**), Quantification of NeuN+ neurons in CA2/3 at 12 months post-injection. CTR treatment significantly increased neuron counts in TDP-43cKO mice compared to GFP controls, though not to the level of control animals. Neuron number in TDP-43cKO-CTR mice approximated the regional CTR infectivity rate (∼30%) (n = 19 (Control), 13 (TDP-43cKO GFP), and 13 (TDP-43cKO CTR) mice. One-way ANOVA, F(2,42) = 48.26, P < 0.0001; Tukey’s post hoc test: Control vs. TDP-43cKO GFP, P < 0.0001; Control vs. TDP-43cKO CTR, P < 0.0001; TDP-43cKO GFP vs. TDP-43cKO CTR, P = 0.0014.). (**D**), Quantification of total hippocampal area based on NeuN-stained sections. Control GFP (n = 18) and Control CTR (n = 18) groups had significantly larger hippocampal areas than TDP-43cKO GFP mice (n = 12) (P < 0.0001). CTR treatment (n = 15) significantly increased hippocampal area in cKO mice compared with GFP-treated cKO animals (P < 0.0001), approaching control group values. (**E**) Immunohistochemical analysis using NeuN antisera in wild-type control (n=20, TDP-43cKo (n=11), TDP-43cKO;Unc CE^-/+^(n=11) and TDP-43cKO;Unc CE^-/-^ (n=11) mice brain (arrow heads indicate degenerated neurons, scale bar 200µm). (**F**) NeuN positive neuron quantification in hippocampal CA2/3 of 4-6 months after TDP-43 deletion. (one-way ANOVA; ns: no significant difference; **P<0.01; ***P<0.005). Scale bars: a: 200 μm, c: 500 μm. Error bars represent mean ± s.e.m. P-values were calculated using one-way ANOVA followed by Tukey’s multiple comparisons test unless otherwise noted. n values and statistical details are provided in Methods.

In TDP-43cKO;*Unc13aCE*^−/−^ mice, hippocampal neuron survival was not preserved. At 6–8 months following tamoxifen-induced TDP-43 deletion, neuron counts in CA2/3 of TDP-43cKO;*Unc13aCE*^−/−^ mice were comparable to those of TDP-43cKO mice and significantly lower than controls (**Fig. 5E-F**). Morphometric analysis showed comparable hippocampal and cortical atrophy across TDP-43cKO and TDP-43cKO;*Unc13aCE*^−/−^ groups (**Sup Fig. 10A-C**), and caspase-3 activation in TDP-43cKO;*Unc13aCE*^−/−^ mice was similar to that of TDP-43cKO mice (**Sup Fig. 10D, right two columns with inset view in lower panel**). Together, these data demonstrate that AAV-PHP.eB-CTR preserves hippocampal neurons and attenuates brain atrophy in TDP-43-deficient mice, whereas such impact did not occur in TDP-43cKO;*Unc13aCE*^−/−^ mice, indicating that synaptic function^50^ and neuronal survival downstream of TDP-43 loss may depend on additional cryptic targets.

## Discussion

A central question for therapeutic development in dementia associated with TDP-43 proteinopathies is whether prevention of cryptic splicing can preserve cognitive function. Using complementary genetic and gene therapy approaches in a conditional TDP-43 knockout mouse model that recapitulates the early TDP-43 dysfunction observed in dementia, we show that genetic ablation of the *Unc13a* cryptic exon is sufficient to preserve recognitive function. AAV-PHP.eB-mediated delivery of CTR rescues the same cognitive deficits and additionally preserves hippocampal neurons and attenuates brain atrophy. Together, these findings establish that prevention of *Unc13a* cryptic splicing is sufficient to preserve cognition downstream of TDP-43 dysfunction and provide *in vivo* preclinical support for therapeutic strategies designed to prevent cryptic splicing in dementia associated with TDP-43 proteinopathy.

AAV-PHP.eB-CTR rescued cognitive deficits to the same extent as abolishing cryptic exon of *Unc13a* but additionally preserved hippocampal neurons and attenuated brain atrophy in TDP-43-deficient mice. By restoring splicing repression across multiple TDP-43 targets (**Fig. 2D-E**), CTR offers an alternative therapeutic strategy that is independent of, and complementary to, target-specific approaches. The autoregulatory element incorporated in CTR further ensures that its expression is upregulated selectively in the context of TDP-43 loss and constrained when endogenous TDP-43 is present^47–49^, a feature that may prove valuable for clinical translation. Long-term CTR exposure in control mice did not reduce hippocampal neuron number or area, nor did it produce overt adverse phenotypes (**Fig. 5C-D**), consistent with this autoregulatory constraint. Companion work in an ALS motor neuron model demonstrates that AAV-PHP.eB-CTR also slows motor neuron disease and prevents paralysis in ChAT-IRES-*Cre;Tardbp*^f/f^ mice^51^, supporting the broad applicability of CTR-based gene therapy across neuronal populations affected by TDP-43 dysfunction.

A key translational challenge is therefore to maximize CTR delivery to affected neurons. In our model, the ∼40% transduction rate of AAV-PHP.eB in hippocampal neurons was sufficient to demonstrate neuroprotection proportional to local transduction efficiency, but clinical translation may require broader coverage. The development of next-generation BBB-crossing AAV capsids with enhanced tropism for human central neurons^52,53^ will be critical to realizing the full therapeutic potential of this approach.

In addition, our finding that *Unc13a* cryptic exon elimination alone is sufficient to rescue cognitive and social deficits in TDP-43-deficient mice provides the first *in vivo* evidence that correcting a single TDP-43 dependent cryptic target can preserve cognition. Our findings resolve ambiguous *in vitro* findings of ASOs targeting *UNC13A* cryptic splicing to restore UNC13A protein levels and synaptic transmission in TDP-43-depleted iPSC-derived neurons^33^ ^50^. Our findings showing sufficiency of targeting solely *Unc13a* cryptic exon to preserve memory encourage clinical testing of ASOs designed to correct *UNC13A* cryptic splicing for dementias associated with TDP-43 proteinopathy. The rescue of recognition memory and social novelty preference by *Unc13a* correction is concordant with the established dependence of these cognitive domains on glutamatergic hippocampal circuits: object recognition memory requires intact hippocampal synaptic function^54^, and social memory is critically dependent on CA2 pyramidal neuron activity^55,56^—the hippocampal subregion most vulnerable to TDP-43-dependent neuron loss in our model. Notably, the impact of Unc13a on behavior is concordant with the human genetic evidence: *UNC13A* harbors the strongest common risk allele for FTLD-TDP^57^, and homozygosity for the risk variant shortens survival by more than three years in FTD patients with TDP-43 pathology^33–35^. Our functional data now offer a mechanistic explanation for this genetic signal: loss of UNC13A-dependent synaptic function upon TDP-43 dysfunction directly undermines the neural circuits required for cognition, and its restoration is sufficient to reverse this circuit-level impairment. These results provide strong preclinical support for the clinical development of UNC13A-directed splice-correcting therapeutics^33–35^ for dementias associated with TDP-43 pathology.

Several limitations should be noted. While both mouse and human harbor TDP-43-dependent cryptic exons targets are mostly different except *UNC13A*, which also reside in different intronic regions, and the extent to which our findings on *Unc13a* correction translate to human UNC13A biology requires further investigation. Additionally, the TDP-43cKO model deletes TDP-43 selectively from forebrain excitatory neurons, recapitulating features of FTLD-TDP, LATE, and AD-LATE, but does not capture the full pathological complexity of these disorders, which include cytoplasmic TDP-43 aggregates, glial dysfunction, and additional cell types not modeled here.

In summary, we provide *in vivo* genetic evidence that prevention of *UNC13A* cryptic splicing is sufficient to preserve cognition in a mammalian model of TDP-43 dysfunction and demonstrate that AAV-PHP.eB-mediated delivery of CTR achieves both cognitive preservation and protection against neuron loss and brain atrophy. These findings provide preclinical support for the clinical development of cryptic splicing-targeting therapeutics, including ASOs and CTR-based gene therapy for human dementia associated with TDP-43 dysfunction.

## Methods

### Animal Models and Genotyping

All animal procedures were approved by the Institutional Animal Care and Use Committee (IACUC) at Johns Hopkins University or the University of Wyoming and conducted in accordance with NIH guidelines. To model TDP-43 loss of function in a cell-type–specific manner, we utilized a previously established conditional Tardbp knockout mouse line (Tardbp^f/f^), in which exon 3 of the Tardbp gene is flanked by LoxP sites^58^(The Jackson Laboratory, stock #017591). We crossed these mice with the *CamkIIa* -CreERT2 driver line to generate a tamoxifen-inducible, excitatory neuron–specific Tardbp knockout model^59,60^ (hereafter referred to as TDP-43cKO).

Experimental cohorts included Tardbp^f/f^; *CamkIIa*-CreERT2 mice (referred to as TDP-43CKO) and Tardbp^f/f^ littermates lacking Cre (referred to as Control). Both sexes were used, and groups were balanced by age and sex. Recombination was induced by feeding mice tamoxifen-containing chow (Envigo, TD.130859, 400 mg/kg) for 4 weeks, followed by a 2-week washout period.

Mice were housed in a 12-hour light/dark cycle with ad libitum access to food and water, and group-housed when possible. Genotyping was performed using DNA extracted from ear biopsies. PCR was conducted using primers specific to the loxP-flanked Tardbp allele and the Camk2a-CreERT2 transgene: PCR products were analyzed by agarose gel electrophoresis. Genotyping Primers are as follows:

*CamKIIa*-Cre F: GACAGGCAGGCCTTCTCTGAA
*CamKIIa*-Cre R: CTTCTCCACACCAGCTGTGGA
*Tardbp* FF F: AACTTCAAGATCTGACACCCTCCCC
*Tardbp* FF R: GGCCCTGGCTCATCAAGAACTG;

The expected product of the *CamkIIa*-Cre primers is 536bp. The expected products of the *Tardbp* FF primers are 376 bp for the floxed product and 230 bp for the wild-type product.

To generate *Unc13a* cryptic exon (CE) deletion mouse model, we used CRISPR technology to generate mice lacking *Unc13a* cryptic exon, we deleted a ∼44 bp region (**Fig. 2a**) from the mouse genome that defined the *<u>Unc</u>13a* <u>c</u>ryptic <u>e</u>xon (termed *Unc CE^-/-^* mice) in house transgenic core facility at Johns Hopkins University. We confirmed the specific deletion of CE sequence (44bp) by amplifying using specific primers aligning outside the CE sequence by PCR using efficient DNA polymerase master mix (**Sup Fig. 3A**) and further by sequencing (**Sup Fig. 3B**).

Specific primers were used to genotype *Unc13a* CE deletion as followed,

Unc13a CE Deletion: F: GGCTCTCCTGTCTTCTCAGC
Unc13a CE Deletion: R: CTTTGTGTGCAAGGCACGAA

To determine the impact of *Unc13a* cryptic exon upon TDP-43 depletion, *UncC^-/-^* mice were crossbred (**Sup Fig. 4B**) with *CaMKII-CreER;Tardbp^f/f^* mice to generate a cohort of mice lacking simultaneously both *Unc13a* cryptic exon and TDP-43 in their cells of TDP-43cKO;UncCE^-/-^ (*CaMKII-CreER;Tardbp^f/f^*;*UncCE^-/-^* mice).

To deplete TDP-43, tamoxifen-containing food administered similarly as explained earlier in this section to mice at the age of 2-months and analyses 6 and 8-months of age for neuropathological and biochemical analysis.

### AAV Vector Design and Delivery

The therapeutic construct CTR (Chimeric TDP-43 Repressor) was designed by fusing the N-terminal RNA recognition domains (RRM1 and RRM2, residues 1–267) of human TDP-43 with the splicing repressor domain of RAVER1 (residues 450–643), replacing the aggregation-prone C-terminal glycine-rich region^1^. The construct includes the endogenous *TARDBP* 3′ untranslated region (3′UTR) to preserve TDP-43 autoregulatory feedback. The CTR transgene was driven by the CBh (chicken β-actin hybrid) promoter.

Control vectors included AAV.PHP-CMV-GFP (Cat# AAV-0090, Virovek), with a reported titer of 2E13 vg/mL, and AAV.PHP-CMV-mCherry, both obtained from the same supplier.

For adult intracerebroventricular (ICV) delivery, mice were anesthetized with isoflurane and placed in a stereotaxic frame. A total of 5 μL (1E11 vg) was injected into the right lateral ventricles at the following coordinates from bregma: AP –0.5 mm, ML 1.0 mm, DV –2.5 mm^61^. Injection was performed at a rate of 1 μL/min using a 5 μL Hamilton syringe. Mice were allowed to recover on a heating pad and monitored until fully ambulatory.

For neonatal injections, pups were injected within 8 h of birth. Cryo-anesthesia was induced by brief exposure to wet ice (<2 min) with a protective barrier to prevent skin injury. A pulled glass capillary needle (41 mm, ∼0.5 mm tip) was inserted ∼3 mm into the lateral ventricle, and 3 μL of AAV9 (1×10^13 vg/mL) encoding either CTR or GFP was slowly delivered over ∼15 s. Trypan blue (0.05%) was included in the injection solution to aid visualization. Following injection, pups were rewarmed under a heat source, placed in bedding from the home cage to restore maternal scent, and returned to the dam once active. Survival was ∼80%, with early losses primarily due to incomplete recovery from anesthesia (P0–P1) and occasional hydrocephalus developing at later stages (P14–P21). Pups showing signs of distress were euthanized according to IACUC guidelines.

### Tissue Collection and Immunohistochemistry

Mice were euthanized at designated timepoints (1-, 3-, 6-, or 12-months post-injection) by isoflurane overdose, followed by transcardial perfusion with phosphate-buffered saline (PBS) and then 4% paraformaldehyde (PFA) in PBS. Brains were carefully dissected and post-fixed in 4% PFA at 4°C overnight, then dehydrated, embedded in paraffin, and sectioned sagittally at 10 μm thickness using a rotary microtome (Leica HistoCore MULTICUT).

For immunohistochemistry, slides were first oven-baked at 60°C for 30 mins, then deparaffinized in xylene and rehydrated through a graded ethanol series (100%, 95%) to water. Antigen retrieval was performed in 10 mM sodium citrate buffer (pH 6.0) by heating slides in a microwave for 4 minutes. After cooling, sections were encircled with a hydrophobic barrier pen and incubated in blocking solution containing 1.5% normal goat serum and 0.1% Triton X-100 in PBS for 1 hour at room temperature. Primary antibodies diluted in blocking buffer were applied overnight at 4°C in a humidified chamber. CTR expression in the mouse was expressed using a N-terminal targeting human TDP-43 antibody. Primary antibodies used:

human-TDP-43 (Ref#WH0023435M1, 1:1000, Millipore Sigma),
mouse-specific TDP-43 (C-Terminus, Ref#12892-1-AP, 1:1000, Proteintech),
Mouse-NeuN (Cat# MAB377, 1:2000, MERCK).
Cleaved Caspase 3 (1:2000; D3E9, Cell signaling technology)

Endogenous peroxidase activity was quenched by incubating slides in 0.3% hydrogen peroxide (H₂O₂) in methanol for 30 minutes, followed by three 10-minute washes in PBST (PBS + 0.1% Tween-20). Slides were incubated with biotinylated secondary antibodies (Vector Laboratories, BP-9100-50, BP-9200-50) for 1 hour at room temperature. Signal was amplified using VECTASTAIN Elite ABC reagent (Vector Laboratories, PK-7100) for 1 hour and visualized by DAB substrate reaction (Vector DAB Peroxidase Substrate Kit, SK4100), monitoring under a microscope for optimal development. Slides were then counterstained with hematoxylin, dehydrated through graded alcohols, cleared in xylene, and mounted with mounting media.

Brightfield images were captured using a ZEISS microscope equipped with a high-resolution digital camera. Quantification was performed using ImageJ with uniform region-of-interest (ROI) definitions applied across experimental groups.

### Cresyl Violet Staining

For cresyl violet (Nissl) staining, deparaffinized and rehydrated sections were immersed in 0.1% cresyl violet acetate solution for 5–10 minutes, differentiated in 95% ethanol, dehydrated through graded alcohols, cleared in xylene, and coverslipped with mounting media. Stained sections were used for morphometric analysis of brain regions.

### Morphometric Analysis

To quantify hippocampal and cortical atrophy, sagittal brain sections stained with cresyl violet or NeuN were imaged using a ZEISS microscope. Hippocampal area, cortical area, cerebellar area, and cortical thickness were measured using ImageJ by manually tracing regions of interest (ROIs) as illustrated in **Sup Fig. 8A**. For each animal, measurements were obtained from three matched sagittal sections, and values were averaged. All measurements were performed by investigators blinded to genotype and treatment group.

### Immunofluorescence Staining

For immunofluorescent staining of mouse brain tissues, to perform immunofluorescence staining on tissue, the paraffin brain sections were deparaffinized and antigen retrieval was performed by boiling for 4 minutes in 10mM sodium citrate buffer to expose the epitope to the antibodies. Nonspecific binding of antibodies was eliminated by incubating with blocking buffer (1.5% normal goat serum in PBS with 0.1% Triton-X) for 1 hour. After blocking, the primary antibodies against the primary antibodies against cleaved caspase 3 (1:1000; D3E9; Cell Signaling Technology). Unbound antibodies were washed out and incubated with secondary conjugated fluorophores (Alexa Fluor 594). The Zeiss Apotome Inverted Fluorescence Microscope (Zeiss, Germany) was used for imaging.

### Immunoblotting

Total protein from the mouse cortex was extracted by homogenization in RIPA buffer. For the analysis of TDP-43 and Unc13a protein expression, total protein was extracted from cortexes of 8-9-months old mice. Protein concentrations were determined, and equal amounts of protein lysates ( 20 μg per lane) were resolved, transferred to polyvinylidene difluoride membranes, and probed with the following antibodies: TDP-43 polyclonal antibody (1:1000; 10782-2-AP, ProteinTech), Unc13a (1:1000; 55053-1-AP, proteinTech), monoclonal anti-β-tubulin III antiserum (1:10,000; T2200, Sigma) and rabbit anti-GAPDH-HRP conjugated antibody (1:5,000; G9545, Sigma). Immunoblots were developed using enhanced chemiluminescence method (Millipore Corp., MA)

### Reverse Transcription PCR (RT-PCR)

Total RNA was extracted from dissected hippocampal tissue using the RNeasy Mini Kit (Qiagen, #74106) according to the manufacturer’s protocol. RNA concentration and purity were assessed using a Nanodrop spectrophotometer (Thermo Fisher, 13-400-519), and RNA integrity was confirmed by agarose gel electrophoresis.

First-strand cDNA was synthesized using the ProtoScript® First Strand cDNA Synthesis Kit (New England Biolabs, E6300) following the manufacturer’s protocol. Each reaction used 500 ng to 1 μg of total RNA and was primed with a mixture of enzyme mix and random primers to ensure coverage of all transcripts.

To assess cryptic exon inclusion, we designed primers flanking known TDP-43–regulated cryptic exons (e.g., *Adnp2, Ap3b2, Bud23, Camk1g, Crem, Ggct, Unc13a, Synj2bp, Tbc1d1, Usp15, Tecpr1,* and *Washc4*)^43^. Products were separated on 1.5% agarose gels and visualized using GelRed staining under UV illumination. Band intensities were quantified using ImageJ and normalized to the total transcript signal (i.e., inclusion + exclusion bands).

To evaluate CTR autoregulation, a primer pair was designed targeting the N-terminal human TDP-43 sequence (RRM1) and the C-terminal RAVER1 fusion domain. Amplification of this junction-specific product confirmed the presence of the CTR transcript.

All PCR reactions were performed in technical duplicates or triplicates, and at least three biological replicates per condition were included. Touch-down thermal cycling conditions were optimized per primer set and are provided in Supplementary Table 2-4.

### BaseScope In Situ Hybridization and RNA–Protein Co-Detection

BaseScope™ in situ hybridization was performed on paraffin-embedded sagittal brain sections (10 μm) using the BaseScope™ v2-RED Detection Kit (ACD BioTechne, Cat# 323910) according to the manufacturer’s protocol. Sections were baked at 60°C for 30 min, deparaffinized through xylene and graded ethanol, and subjected to target retrieval and protease digestion as specified. Probe sets targeting the Unc13a (Cat# 1182491-C1), Synj2bp (Cat# 712191) cryptic exons, and CTR mRNA (Cat# 1573691-C1), were hybridized for 2 hr at 40°C, followed by sequential AMP amplification and RED chromogenic development. Slides were counterstained with 50% hematoxylin and mounted with AquaMount.

For simultaneous protein detection, a subset of sections underwent RNA–protein co-detection using the RNA-Protein Co-Detection Ancillary Kit (Cat# 323180) following the manufacturer’s protocol (MK 51-149/Rev B). Briefly, target retrieval was performed in 1× Co-Detection Target Retrieval solution at 98–102°C for 15 min; primary antibody diluted in Co-Detection Antibody Diluent was applied overnight at 4°C; and sections were post-fixed in 10% neutral buffered formalin for 30 min before protease digestion and probe hybridization as above. After RED chromogenic development, IHC detection was completed using Co-Detection Blocker, secondary antibody, ABC reagent, and DAB. Sections were blued in 0.02% ammonia water and mounted after xylene clearing.

Brightfield images were acquired on a ZEISS microscope. BaseScope™ puncta per nucleus were counted manually within defined hippocampal regions of interest using ImageJ, with at least three sections per mouse and 3–5 mice per group analyzed under blinded conditions.

### Behavioral Testing

Behavioral experiments were conducted at the University of Wyoming Animal Behavior Core Facility under blinded conditions. All testing was performed during the light phase (9:00 a.m. to 5:00 p.m.), and mice were habituated to the testing room for at least 30 minutes before each assay.

Open field testing was used to evaluate general locomotion and anxiety-like behavior. Mice were placed in a 42 × 42 cm open-field arena for 15 minutes. Distance traveled and time spent in the center zone were tracked using Noldus EthovisionXT 15 software.

Novel object recognition (NOR) was used to assess recognition memory. Mice were exposed to a 42 × 42 cm open-field arena settled with two identical objects for 10 minutes. After a 5-minute delay, the arena was cleaned and one object was replaced with a novel object of similar size, and mice were reintroduced to the arena for 10 minutes. Object exploration was scored manually by trained observers blinded to group allocation.

Social behavior testing was performed and recorded in an adapted same-chambered procedure as previously described^62^. During habituation, mice were allowed to freely explore a 42 x 42 cm arena settled with two empty wire cages at the opposite corners for 10 minutes. In the 10-minute sociability test phase, an unfamiliar age- and sex-matched conspecific (stranger 1) was enclosed in one of the empty wire cages. In the 10-minute social novelty test phase, a second unfamiliar age- and sex-matched conspecific (stranger 2) was placed in the previously empty wire cage. Time spent sniffing or touching each cage was scored manually frame by frame by trained observers blinded to group allocation.

All behavioral assessments were performed by experimenters blinded to genotype and treatment group. Data were analyzed as described in the statistical analysis section.

### *In vivo* calcium imaging

*In vivo* calcium imaging was performed to monitor neuronal activity in the prelimbic cortex of TDP-43cKO and control mice. 500nl of AAV1-CamKII-GCaMP6f (Addgene, 2 E13 GC/mL) was injected stereotaxically into the prelimbic cortex (coordinates: A/P +1.9 mm, M/L 0.5 mm, D/V 1.75 mm) under isoflurane anesthesia as previously described^63^. A 1 mm diameter GRIN lens (Grintech) was implanted into the injection site to reach the depth of 1.8 mm and secured with dental cement as previously described^63^. Mice were allowed to recover for at least 4 weeks prior to recording.

*In vivo* Ca^2+^ Imaging was performed using a previously established custom-built miniature fluorescence microscope recording system^64^ during exploratory behavior in a 42 x 42 cm open-field arena. Three recording sessions, each lasting 5 minutes per mouse, were conducted. Calcium fluorescence signals were acquired at 10 Hz and preprocessed using standard motion correction and ΔF/F0 normalization. Individual neurons were identified and segmented using CNMF-E implemented in MATLAB. Calcium event rates were quantified by thresholding deconvolved signals, and the average spike rate per neuron was calculated for each animal as previously described^65^.

Mice were excluded from analysis if recordings lacked sufficient signal-to-noise or spatially stable fields of view. Of 67 mice that underwent imaging procedures, 30 (45%) produced data of sufficient quality for analysis (see Supplementary Table 1 for group breakdown). Sample sizes per group ranged from 5 to 9 animals. All animals were included in downstream group-level comparisons.

### Data analysis

All statistical analyses were performed using GraphPad Prism (version 10) unless otherwise specified. Data are presented as mean ± standard error of the mean (s.e.m.), unless otherwise indicated. Group comparisons were evaluated using unpaired two-tailed Student’s t-tests, Mann-Whitney U test, one-way ANOVA, or two-way ANOVA with Holm–Sidak post hoc correction, depending on experimental design and variance structure.

## Acknowledgements

This work was supported in part by the National Institutes of Health grants R01 NS095969 (to P.C.W.), UG3/UH3 NS115608 (to P.C.W.), R33NS115161 (to P.C.W.), R01 NS129878 (to P.C.W. and Y.L.), and the Intramural Research Program of the National Institutes of Health (NIH). The contributions of the NIH author (D.-T.L.) are considered Works of the United States Government. Acknowledgements to Eliya Karoutchy, Jiajun Qiu, Kerry Xiaoke Chen, and Nancy Yan, who provided technical help. The findings and conclusions presented in this paper are those of the author(s) and do not necessarily reflect the views of the NIH or the U.S. Department of Health and Human Services.

## Author contributions

T.C., M.S.B and P.C.W. conceptualized, designed, and interpreted the study. T.C., Y.L., M.S.B., J.P.L., and P.C.W. wrote the manuscript. T.C., R.T., R.L., A.P.M., I.R.S., G.D.B., B.P., C.J., and X.W. performed experiments. D.-T.L. provided the custom-built miniscope recording system. All the authors reviewed and approved the final manuscript.

## Supplementary data figure legends

**Supplementary Figure 1.**
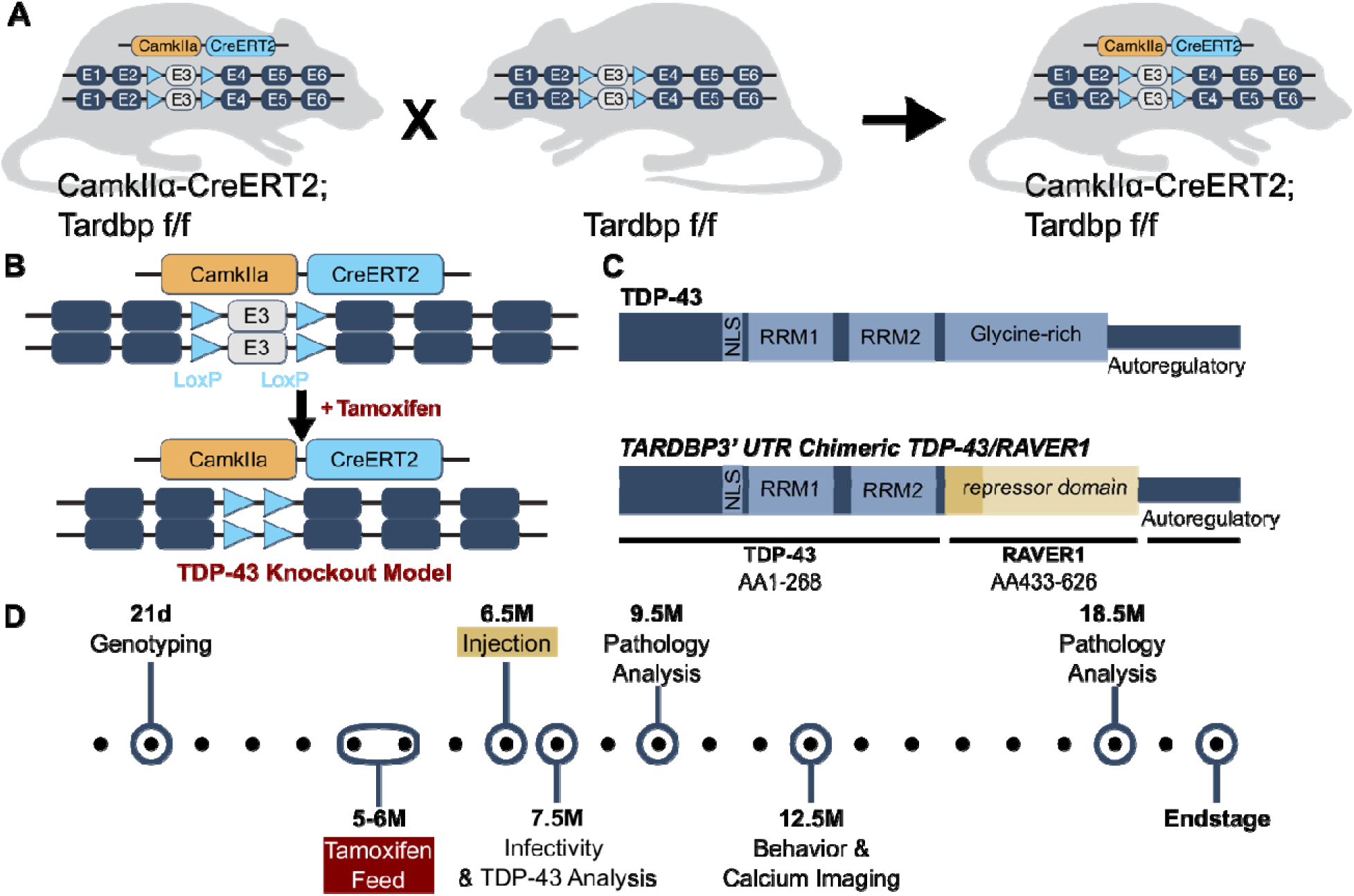
Conditional TDP-43 knockout model and AAV-CTR vecto therapeutic design. (**A, B**), Schematic of the inducible, excitatory neuron–specific *Tardbp* conditional knockout (TDP-43CKO) mouse model. LoxP sites flank exon 3 of *Tardbp* in *Tardbp*^f/f^ mice, and CreERT2 is driven by the Camk2a promoter. Upon tamoxifen administration, exon 3 is excised in forebrain excitatory neurons, resulting in loss of functional TDP-43. (**C**), Diagram of the CTR construct (Chimeric TDP-43 Repressor), composed of the N-terminal RNA recognition motifs (RRM1 and RRM2) of TDP-43 (amino acids 1–267), fused to the splicing repression domain (RAVER1, amino acids 450–643), and followed by the endogenous human *TARDBP* 3′ untranslated region (3′UTR) to preserve autoregulatory feedback. (**D**) Timeline of the *in vivo* study design. Mice were fed tamoxifen at 5 months of age for 1 month to induce recombination. After a 2-week recovery, intracerebroventricular injection of AAV-PHP.eB vectors expressing CTR or GFP were used. Mice were analyzed at 1-, 3-, 6-, and 12-month post-injection.

**Supplementary Figure 2.**
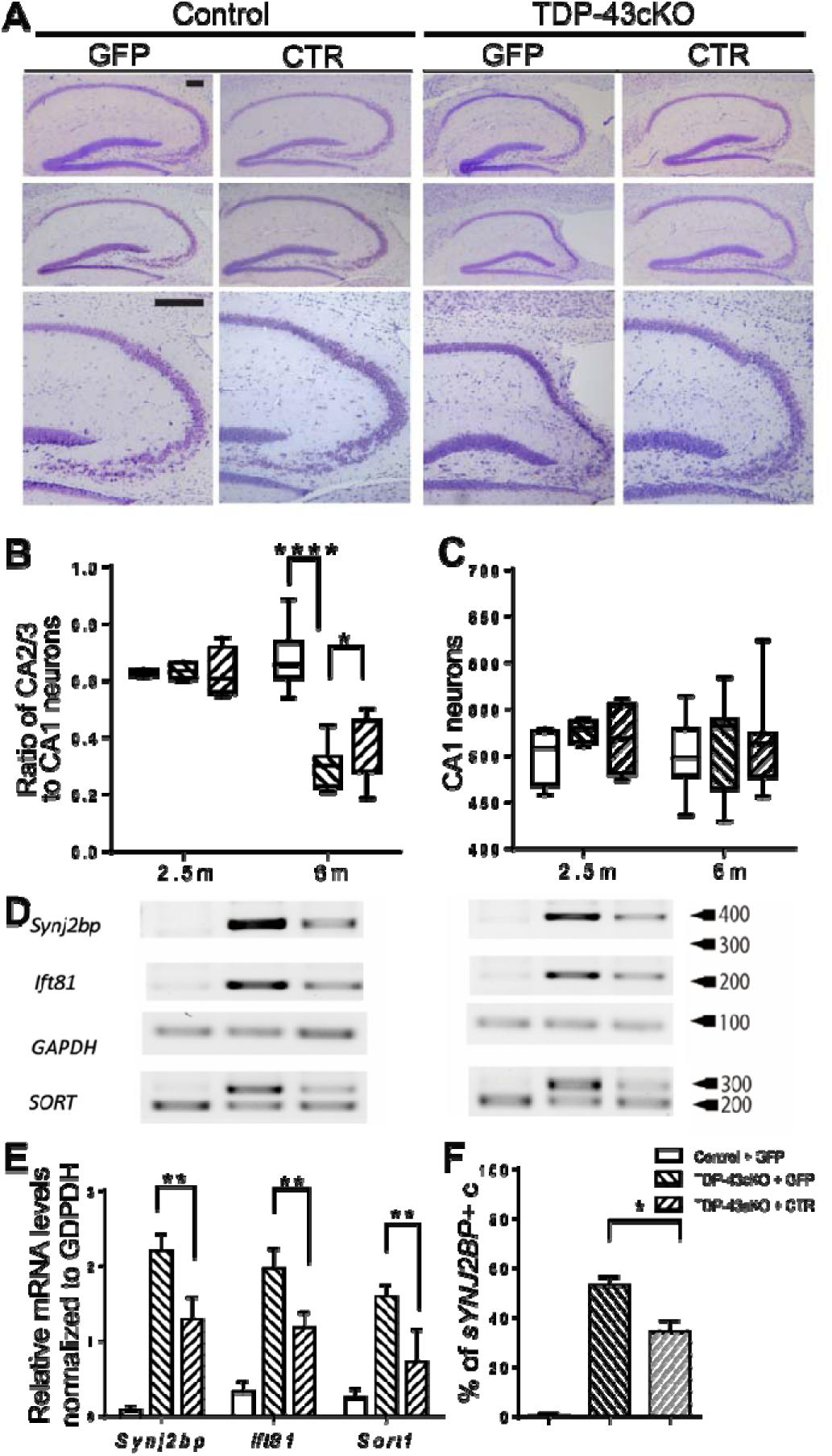
AAV9-ICV Delivery of CTR at P0 Attenuates Cryptic Exon Inclusion of TDP-43 Target Genes and Preserves Neuronal Integrity in the Hippocampus. **(A)** Representative cresyl violet staining of the hippocampus in Control + GFP, Control + CTR, TDP-43cKO + GFP, and TDP-43cKO + CTR mice at 2.5 and 6 months of age. **(B),** Quantification of the ratio of neurons in the CA2/3 region, normalized to neuron numbers in CA1 region, 2.5 m time point: Control + GFP (n = 4), TDP-43cKO + GFP (n = 4), and TDP-43cKO + CTR (n = 4), 6 m time point: Control + GFP (n = 9), TDP-43cKO + GFP (n = 9), and TDP-43cKO + CTR (n = 12) **(C)**, Quantification of neurons in the CA1 region in the same groups as panel B. **(D)** Representative semi-quantitative RT-PCR analysis of three TDP-43 target cryptic exon RNAs (Synj2bp, Ift81, and Sort) in hippocampus and cortex of Control + GFP (n = 3), TDP-43cKO + GFP (n = 4), and TDP-43cKO + CTR (n = 3) mice at 4 months of age. **(E)**, Quantification of cryptic Synj2bp, Ift81, and Sort RT-PCR cryptic products in hippocampus from the same groups as in (d). **(F)**, Quantification of cryptic Synj2bp positive cells by *in situ* hybridization in the hippocampus from the same groups. Band intensities were quantified using ImageJ. One-way ANOVA: **** P < 0.0001, ** P < 0.01, * P<0.05. Scale bars: 200um.

**Supplementary figure 3.**
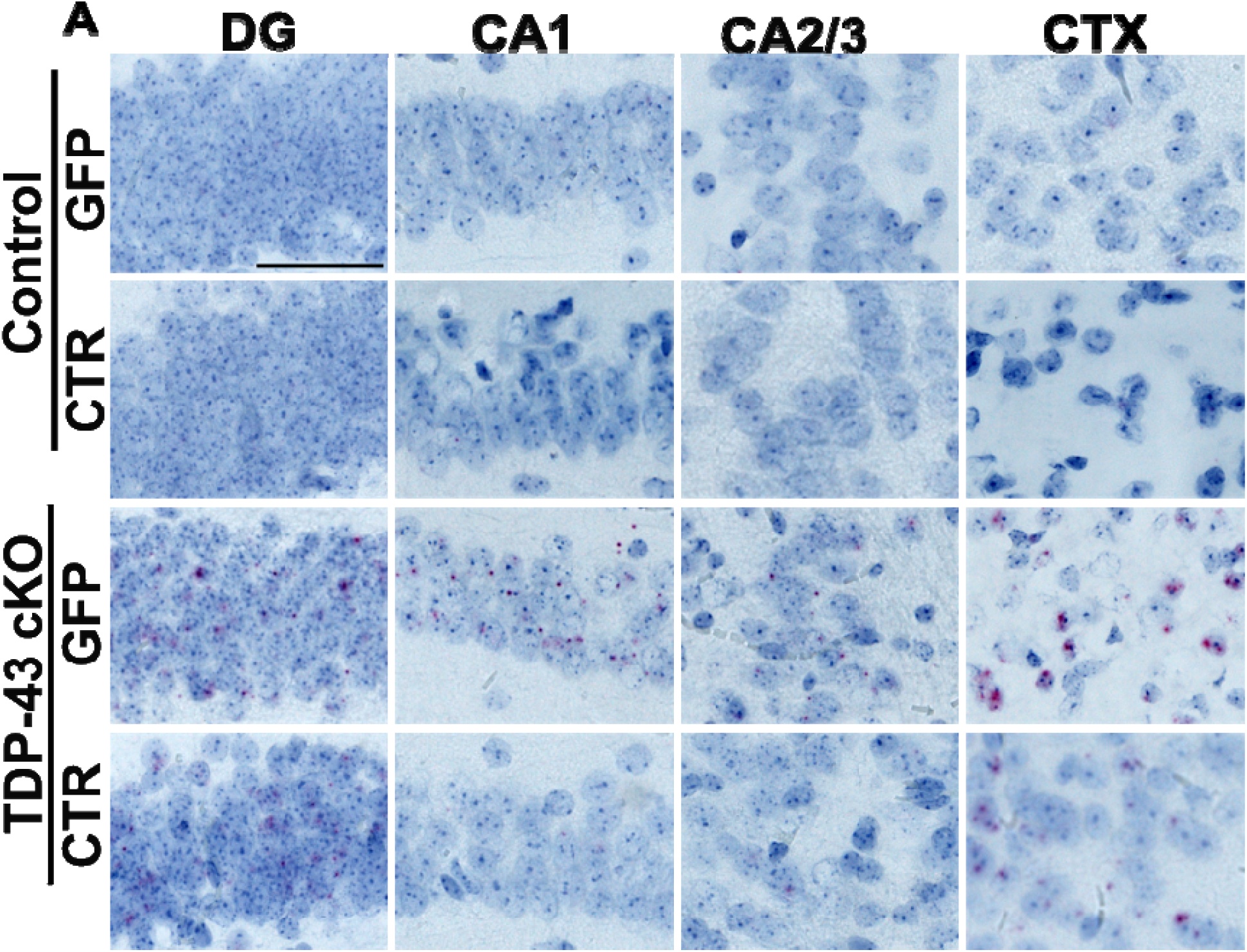
CTR restores cryptic *Unc13a* repression in TDP-43cKO and repressed in control mice. (**A**) Representative images of BaseScope *in situ* hybridization for cryptic *Unc13a* RNA reveal the spatial distribution of splicing defects across hippocampal subregions. Cryptic *Unc13a* transcripts were elevated in dentate gyrus (DG), CA1, and CA2/3 of TDP-43cKO-GFP mice and markedly reduced in CTR-treated animals, indicative of CTR restoring TDP-43 splicing regulation. Scale bars a: 100 μm.

**Supplementary figure 4.**
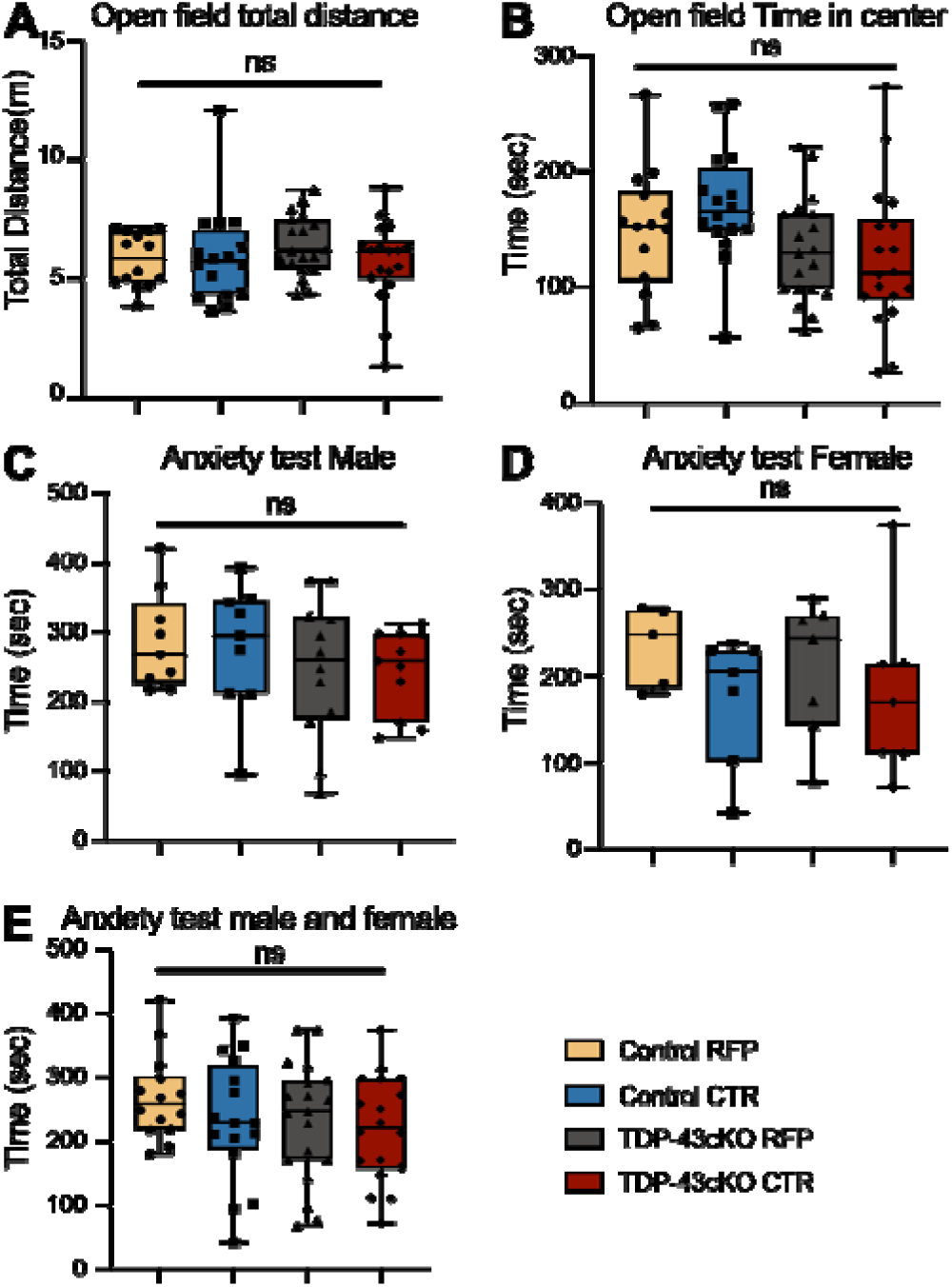
No significant differences were observed in locomotor activity or anxiety-related behavior between genotypes or treatment groups in the open field and light–dark box tests. **(A)** Total distance traveled in the open field test. **(B)** Time spent in the center of the open field arena. **(C)** Time spent in the light compartment of the light–dark box test in male mice. **(D)** Time spent in the center of the open field in female mice. **(E)** Time spent in the light compartment of the light–dark box test in male and female mice combined. Groups: Control + RFP(N = 16), Control + CTR(N = 14), TDP-43cKO + RFP(N = 18), and TDP-43cKO + CTR(N = 19). Data are presented as mean ± SEM; ns = not significant by one-way ANOVA.

**Supplementary figure 5:**
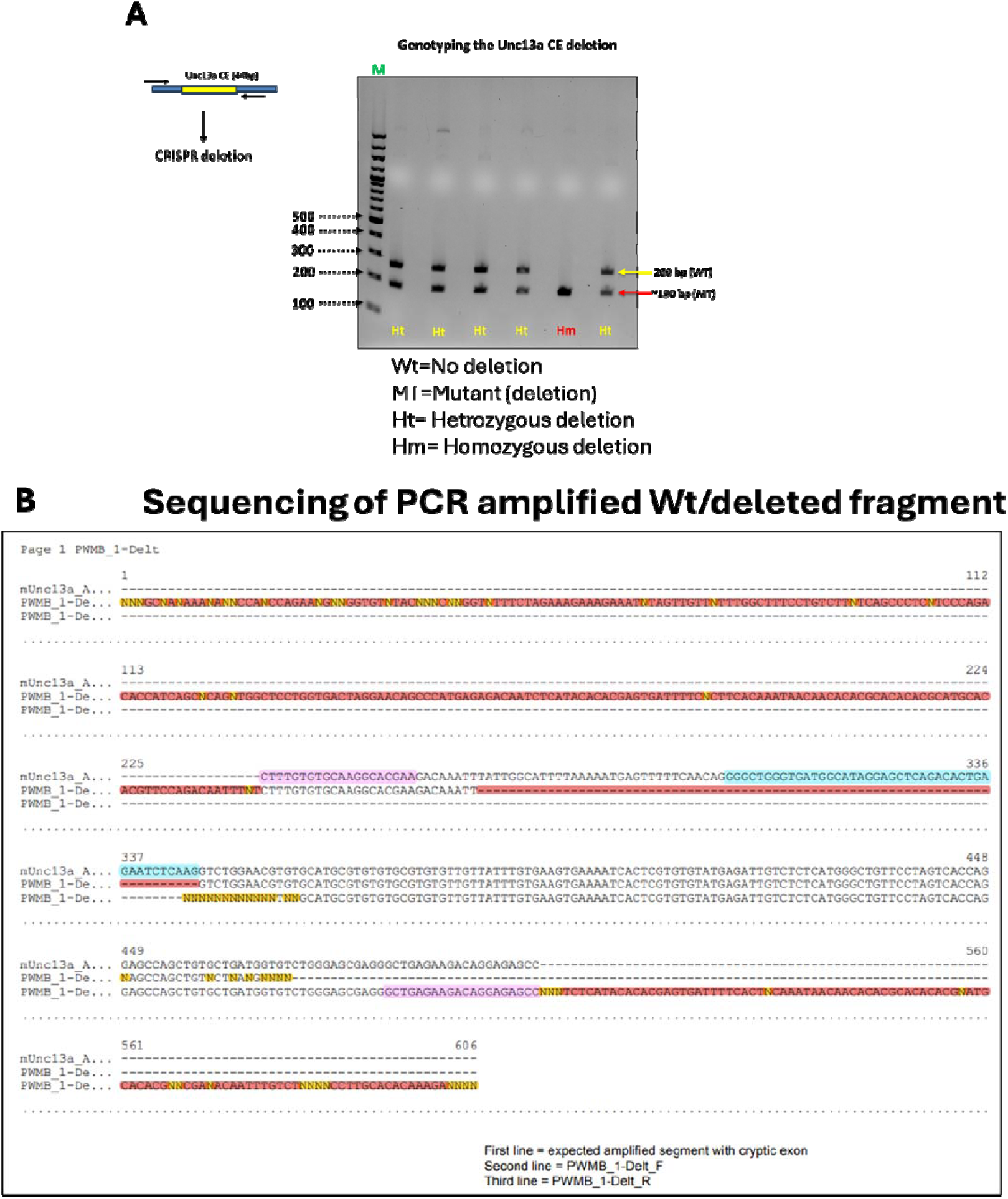
CRISPR deletion of Unc13a CE sequence from mouse genome. (**a**) Schematic presentation of *Unc13a* CE sequence (44bps) was deleted by CRISPR technology, specific primer set binds out side the cryptic sequence to genotype deletion (∼260bp, wild-type (WT) fragment; ∼190 bp, mutant or deleted (MT). Different lanes showing individual mouse having Hetrozygous and Homozygous genotypes for *Unc13a* CE deletion. (**b**) Wild-type and deleted fragments were sequences and align with sequence database, first line: expected amplified segment with cryptic sequence highlighted in cyan blue; second line: PWMB_1-Delt_F is forward primer amplified segment without cryptic sequence highlighted in orange; third line: PWMB_1Delt_R is reverse primer amplified segment without cryptic sequence. Forward and reverse primer sequence are highlighted in pink color.

**Supplementary figure 6.**
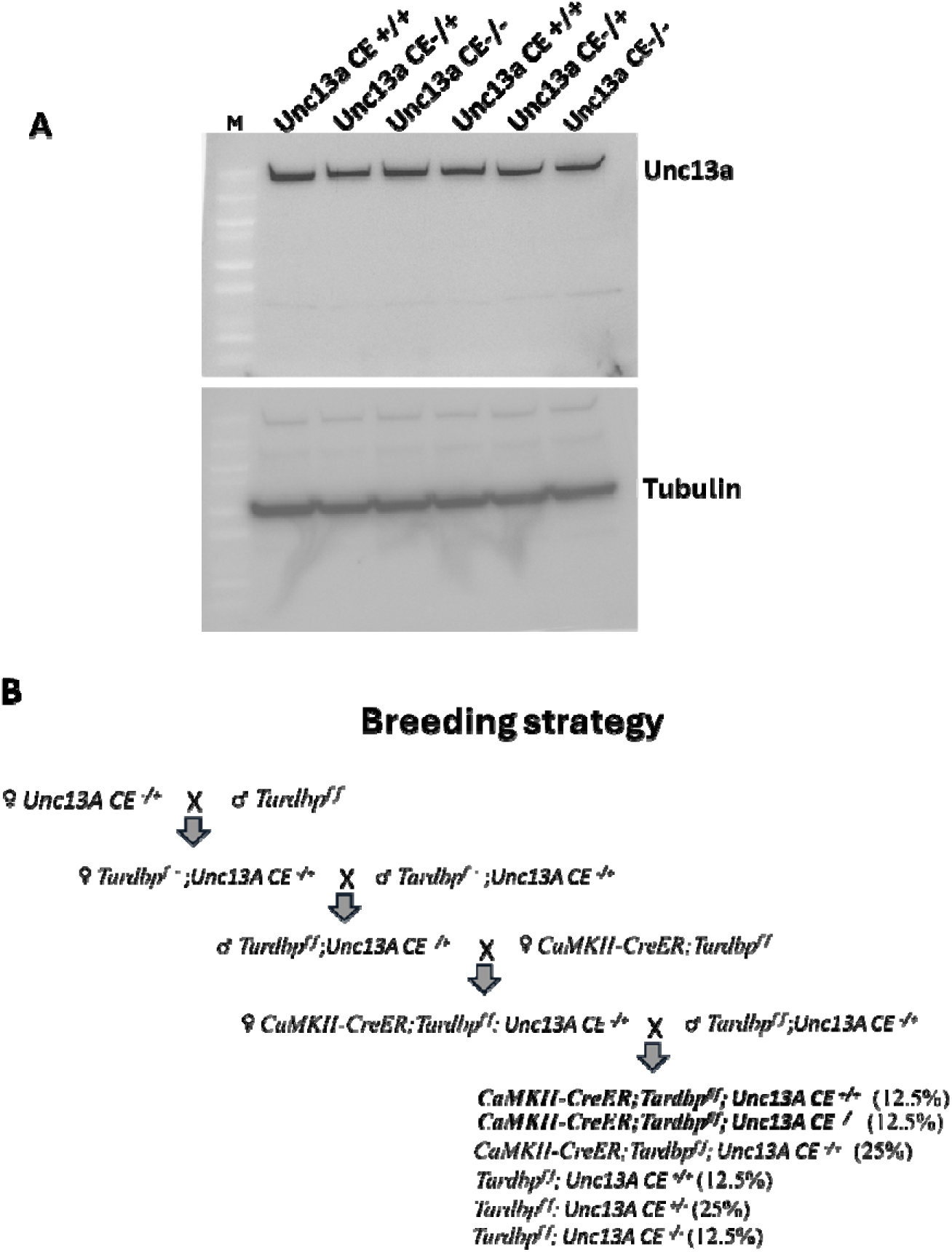
Deletion of the intronic Unc13a CE sequence itself has no impact on its protein expression. **(a)** Immunoblotting analysis of Unc13A protein in wild-type and Unc13a CE deleted mice showing no change in protein expression**. (b)** Breeding strategy for crossbreeding Unc13a CE deleted mouse with CaMKII-CreER;Tardbpf/f mice to produce CaMKII-Cre;Tardbpf/f;UncCE^-/-^ including respective littermates.

**Supplementary figure 7.**
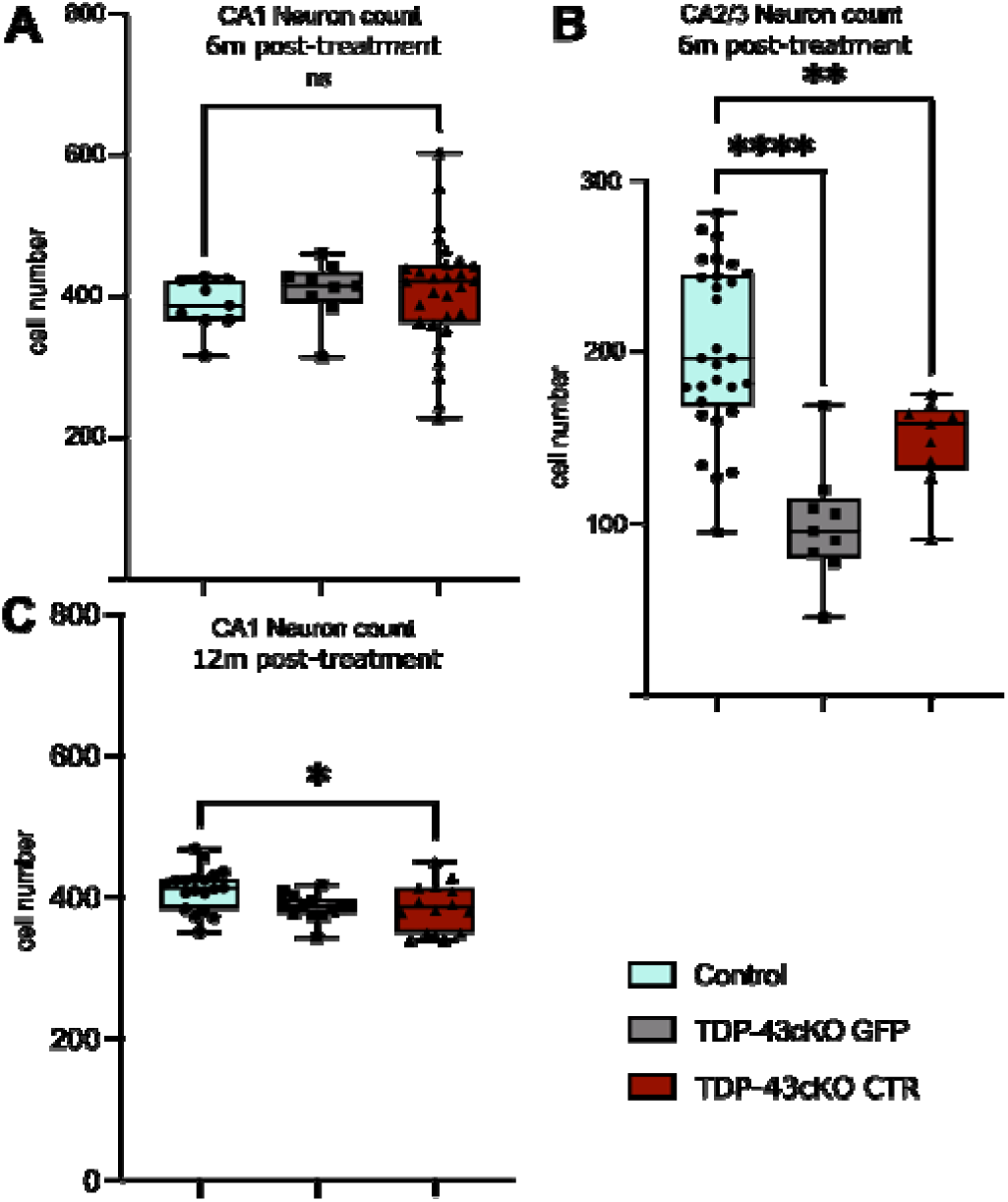
Neuronal Survival after CTR Treatment. (**A**) Quantification of CA1 neuron numbers at 6 months post-treatment in Control(n=29), TDP-43cKO + GFP(n=9), and TDP-43cKO + CTR(n=9) mice. No significant differences were observed between groups. (**B**) Quantification of CA2/3 neuron numbers at 6 months post-treatment in the same groups as in (a). TDP-43cKO + GFP mice showed a significant reduction compared to Control, and CTR expression partially rescued neuron numbers. (**C**) Quantification of CA1 neuron numbers at 12 months post-treatment in in Control(n=19), TDP-43cKO + GFP(n=13), and TDP-43cKO + CTR(n=13) mice. Data are presented as mean ± SD. Statistical analysis was performed using one-way ANOVA followed by post hoc multiple comparisons. Statistical significance: p < 0.05 (*), p < 0.01 (**), p < 0.0001 (***), ns = not significant.

**Supplementary figure 8.**
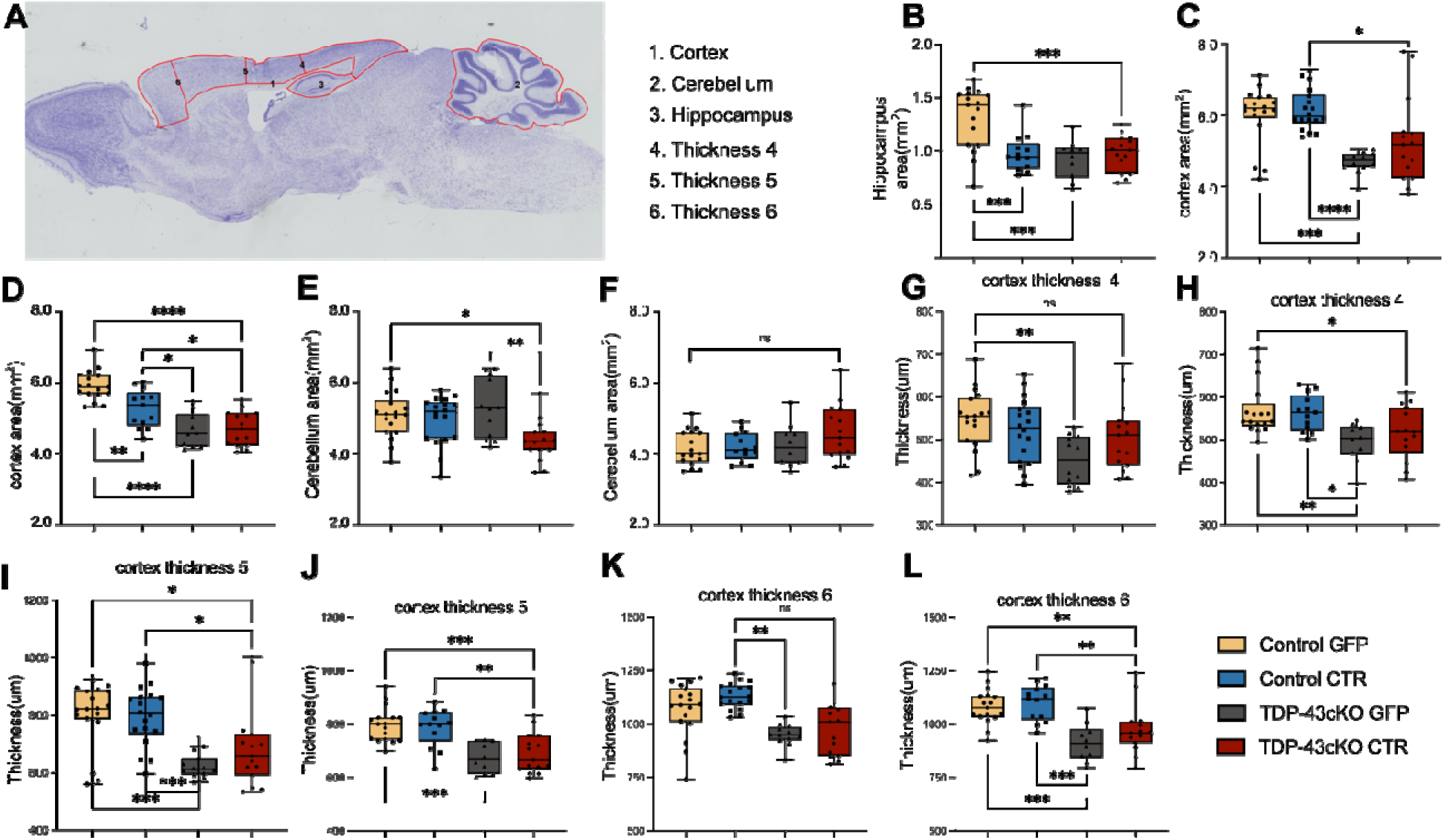
Morphometric analysis of cortical thickness, cerebellar area, and hippocampal area after CTR treatment in Control and TDP-43cKO mice. (**A**) Schematic illustration of the brain regions measured for cortical thickness and area, cerebellar area, and hippocampal area. Red outlines indicate the regions of interest (ROIs): cortex (1, 4–6), cerebellum (2), and hippocampus (3). (**G, I, K**) Quantification of cortical thickness in the indicated cortical regions at 6 months after treatment across experimental groups: Control + GFP(n=10), Control + CTR(n=15), TDP-43cKO + GFP(n=16), and TDP-43cKO + CTR(n=13). (**H, J, L**) Quantification of cortical thickness in the indicated cortical regions at 12 months after treatment across experimental groups: Control + GFP(n=18), Control + CTR(n=18), TDP-43cKO + GFP(n=12), and TDP-43cKO + CTR(n=15). (**B, D, F**) Quantification of cerebellar area (F), cortical area (D), and hippocampal area (B) at 6 months after treatment. (**C, E**) Quantification of cerebellar area (E) and cortical area (C) at 12 months after treatment. Data are presented as mean ± SD. Statistical analysis was performed using one-way ANOVA followed by post hoc multiple comparisons. Statistical significance: p < 0.05 (*), p < 0.01 (**), p < 0.0001 (***), ns = not significant.

**Supplementary Figure 9.**
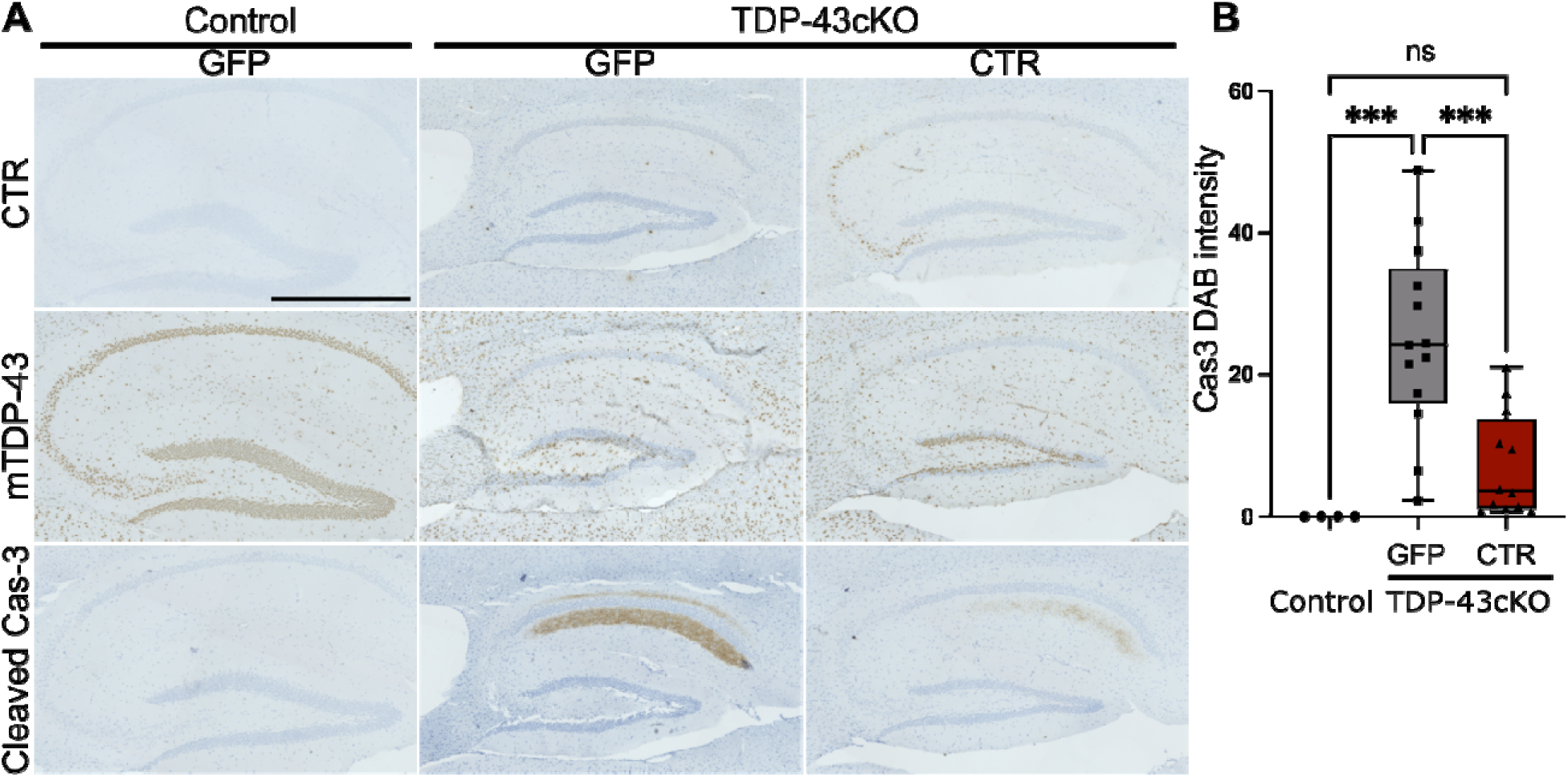
AAV-PHP.eB-CTR attenuates caspase-3 activation in TDP-43cKO mice. (**A**) Immunohistochemistry of the hippocampus showing CTR, mouse TDP-43, and Cleaved Caspase-3. Robust caspase-3 activation was observed in TDP-43cKO mice treated with GFP, whereas CTR-treated cKO mice displayed markedly reduced activation. (**B**), Quantification of caspase-3 DAB intensity in the hippocampus at 12 months post-injection. GFP-treated TDP-43cKO mice (N=4) showed significantly higher caspase-3 staining compared to CTR treated TDP-43cKO mice (N=12) and GFP-treated control mice (N=13). CTR treatment rescued caspase-3 activation in TDP-43cKO mice to levels indistinguishable from controls. One-way ANOVA, F(2,26) = 13.78, P < 0.0001; Tukey’s post hoc test: Control GFP vs. TDP-43cKO GFP, P=0.0007; Control GFP vs. TDP-43cKO CTR, P=0.4693; TDP-43cKO GFP vs. TDP-43cKO CTR, P = 0.0005.). Scale bars: a: 500 μm. Error bars represent mean ± s.e.m.

**Supplementary Figure 10.**
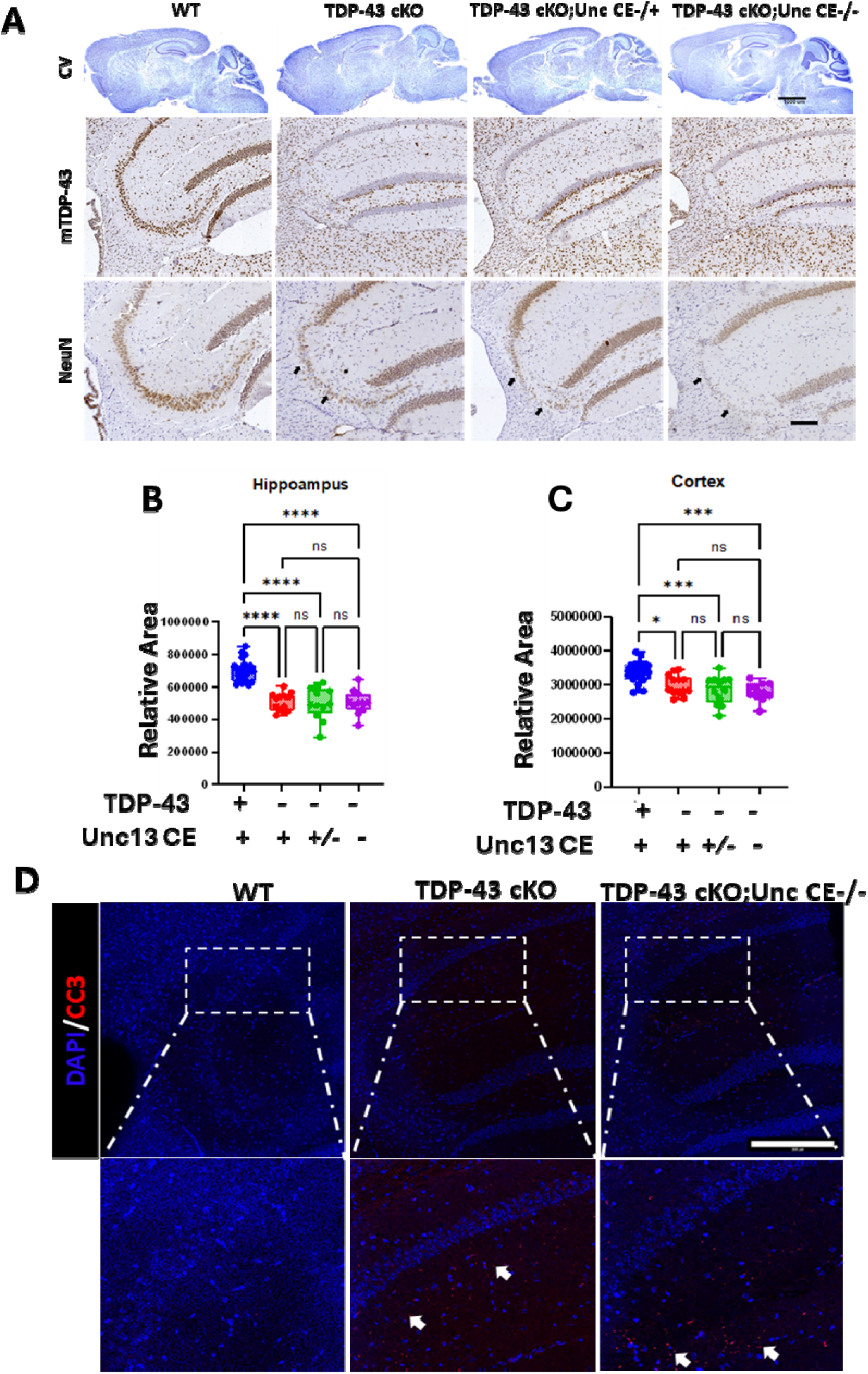
Prevention of *Unc13a* CE have no rescue effect on neuron loss and brain atrophy in TDP-43 lacking mice. **(a)** Histopathological analaysis of wild-type control (n=15), TDP-43cKO (n=8), TDP-43cKO;UncCE^-/+^ (n=11) and TDP-43cKO;UcnCE^-/-^ (n=11) mice, upper panel showing CV (scale bar, 1000µm), immunohistochemistry of mouse TDP-43 protein (middle panel) and NeuN (lower panel) (scalebar, 200µm). **(b-c)** Measurement of relative areas of hippocampus and cortex of wild-type control (n=15), TDP-43cKO (n=8), TDP-43cKO;UncCE^-/+^ (n=11) and TDP-43cKO;UcnCE^-/-^ (n=11) (one-way ANOVA; ns: no significant difference; **P<0.01; ***P<0.001). **(d)** Immunofluorescence analysis of cleaved caspase 3 (CC3) in wild-type control, TDP-43cKO and TDP-43cKO;UcnCE-/-, showing Unc13a CE deletion could not prevent CC3 levels in TDP-43 deficient mice (arrowhead indicates CC3 positive puncta around hippocampal CA2/3 area in inset view, scale bar, 200µm).

## Supplementary Tables

**Supplementary Table 1.** Perioperative mortality and calcium imaging data by sex and experimental group.

|  | Male(n =<br>41) | Female(n<br>= 26) | Total(n =<br>67) |
| --- | --- | --- | --- |
| Deaths during/after surgery, n (%) | 7 (17%) | 5 (19%) | 12 (18%) |
| Mice with high-quality calcium imaging data, n (%) | 13 (32%) | 17 (65%) | 30 (45%) |
| Breakdown of mice with high-quality data |  |  |  |
| Control + CTR | 3 | 4 | 7 |
| Control + RFP | 3 | 3 | 6 |
| TDP-43cKO + CTR | 2 | 4 | 6 |
| TDP-43cKO + RFP | 5 | 6 | 11 |

**Supplementary Table 2.** RT-PCR protocol using touchdown PCR.

| S<br>tage | Temperature | T<br>ime | Cy<br>cles |
| --- | --- | --- | --- |
| 1 | 95 | 1:00 | 1 |
| 2 | 95 | 0:30 | 10 |
|  | 64, descend 1 degree per cycle | 0:15 |  |
|  | 72 | 0:15 |  |
| 3 | 95 | 0:30 | 30 |
|  | 53 | 0:15 |  |
|  | 72 | 0:15 |  |
| 4 | 72 | 7:00 | 1 |
| Fi<br>nal | 4 | ∞ |  |

**Supplementary Table 3.**
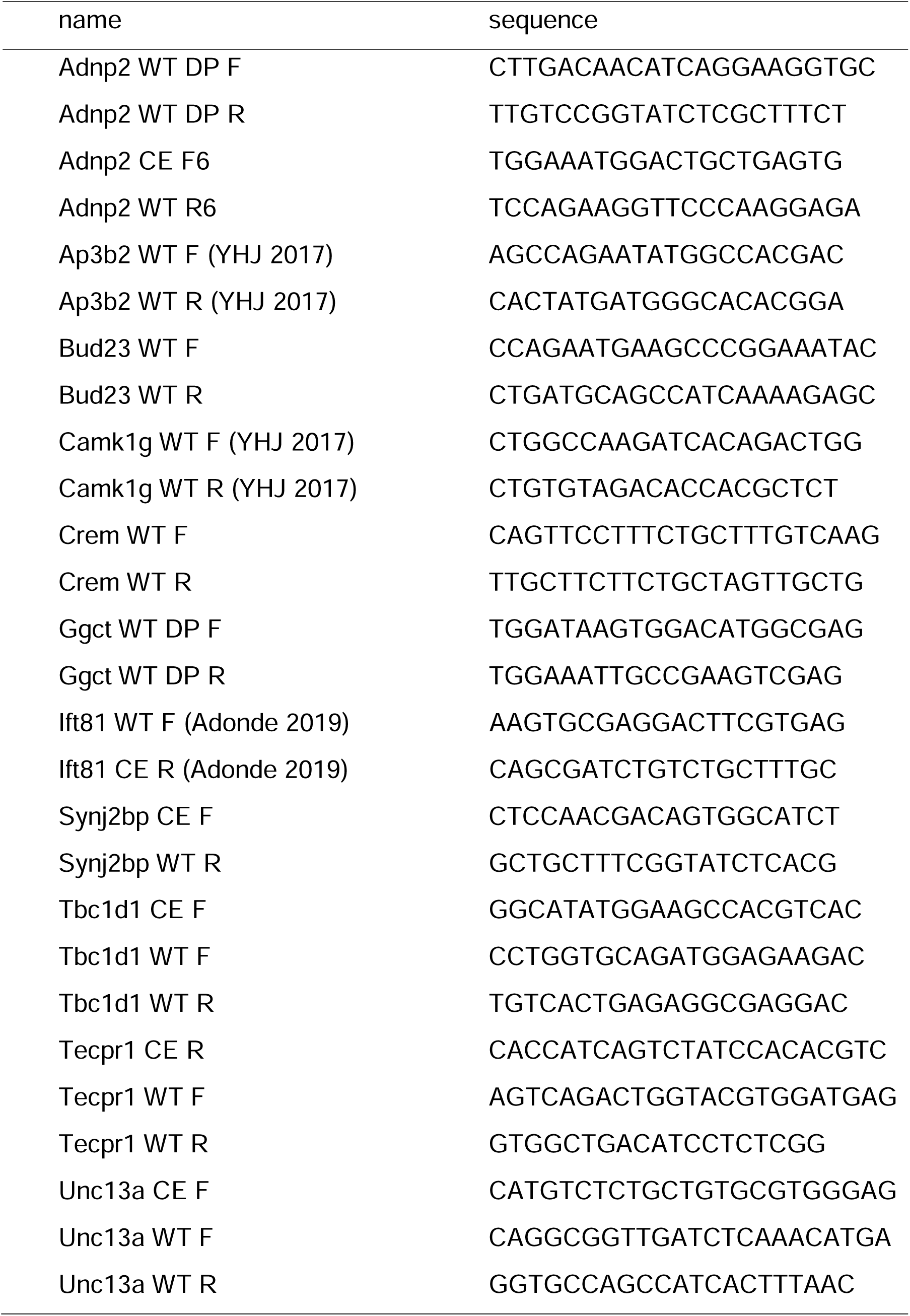
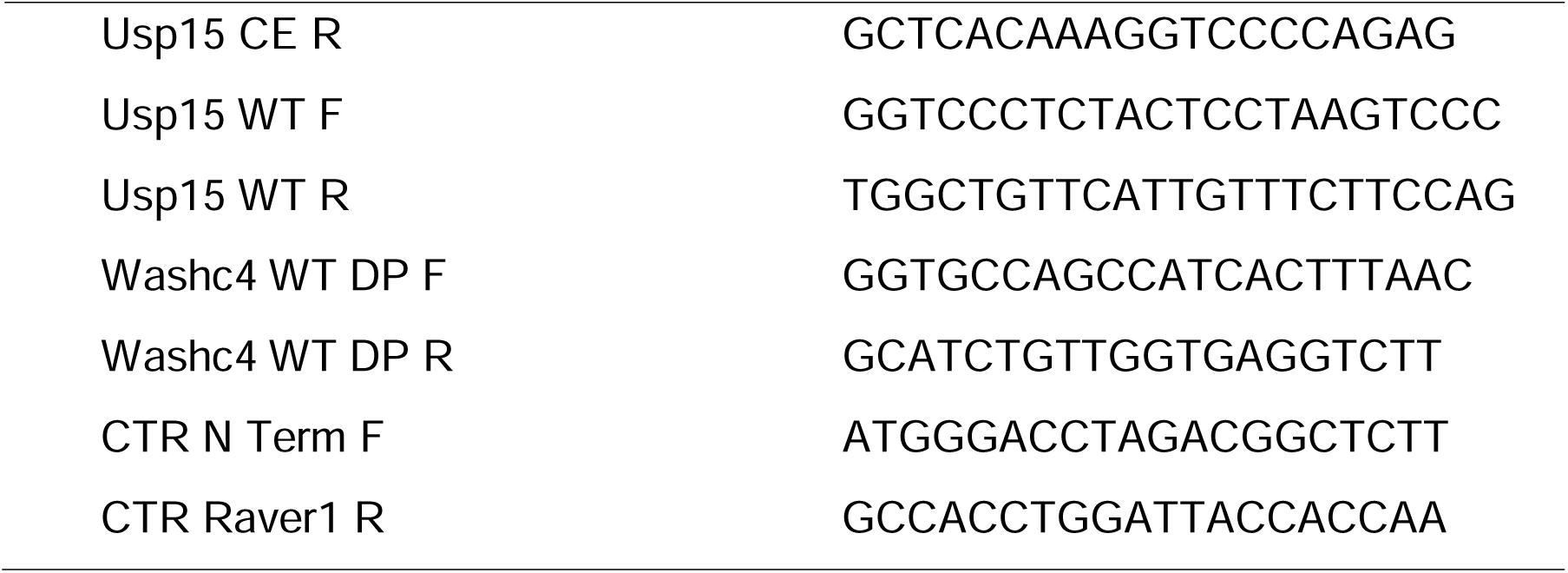
RT-PCR Primers.

**Supplementary Table 4.** Expected RT-PCR band sizes.

| Target | WT band | CE band | Single Band |
| --- | --- | --- | --- |
| Adnp2 | 187 | 339 | 128 |
| Ap3b2 | 374 | 473 | N/A |
| Bud23 | 306 | 409 | N/A |
| Camk1g | 423 | 509 | N/A |
| Crem | 332 | 476 | N/A |
| Ggct | 183 | 285 | N/A |
| Ift81 | N/A | N/A | 187 |
| Synj2bp | N/A | N/A | 326 |
| tbc1d1 | 128 | 238 | 197 |
| Tecpr1 | 329 | 382 | 358 |
| Unc13a | 169 | 213 | 166 |
| usp15 | 192 | 356 | 212 |
| Washc4 | 274 | 472 | N/A |
| CTR | N/A | N/A | 345 |

